# Male mammalian meiosis and spermiogenesis is critically dependent on the shared functions of the katanins KATNA1 and KATNAL1

**DOI:** 10.1101/2022.11.11.516072

**Authors:** Jessica EM Dunleavy, Maddison Graffeo, Kathryn Wozniak, Anne E O’Connor, D. Jo Merriner, Joseph Nguyen, Ralf B Schittenhelm, Brendan J Houston, Moira K O’Bryan

## Abstract

Katanin microtubule severing enzymes are potent M-phase regulators in oocytes and somatic cells. How the complex, and evolutionarily critical, male mammalian meiotic spindle is sculpted remains unknown. Here, using multiple single and double gene knockout mice, we reveal that the canonical katanin A-subunit, KATNA1, and its close paralogue, KATNAL1, together execute multiple aspects of meiosis. We show KATNA1 and KATNAL1 collectively regulate the male meiotic spindle, cytokinesis and midbody abscission, in addition to diverse spermatid remodelling events, including Golgi organisation, and acrosome and manchette formation. We also define KATNAL1-specific roles in sperm flagella development, manchette regulation, and sperm-epithelial disengagement. Finally, using proteomic approaches we define the KATNA1, KATNAL1, and KATNB1 mammalian testis interactome, which includes a network of cytoskeletal and vesicle trafficking proteins. Collectively, we reveal the presence of multiple katanin A-subunit paralogs in mammalian spermatogenesis allows for ‘customized cutting’ via neofunctionalization and protective buffering via gene redundancy.

## Introduction

The katanins are an evolutionary conserved family of microtubule (MT) severing proteins that were first identified as an essential meiosis regulator in *Caenorhabditis elegans* oocytes^1^. Since then, they have been established as a nuanced regulator of MTs, able to employ the same action (physical removal of tubulin heterodimers from the MT lattice) to amplify, disassemble or even strengthen MTs (for reviews see McNally et al.^2^ and Sarbanes et al.^3^). At a cellular level these actions drive rapid MT remodelling events required in many processes including ciliogenesis, cell division and differentiation, and allow fine tuning of homeostatic MT structures and processes.

The canonical katanin complex consists of an AAA ATPase catalytic severing enzyme KATNA1 (aka p60 or A-subunit) and a WD40-repeat containing regulatory subunit KATNB1 (aka p80 or B-subunit) and is conserved across most eukaryotes. In its active ATP bound state, the A-subunit forms a spiral hexamer which binds MTs and threads the C-terminal tail (CTT) of a tubulin subunit through its central pore. An ATP-driven conformational change in the hexamer, causes an extractive force on the tubulin CTT removing the tubulin heterodimer from the lattice. In higher eukaryotes, the B-subunit is not essential for A-subunit severing activity, however it increases MT affinity and severing efficiency, and at a cellular level is often essential for A-subunit localisation^2^. In vertebrates, *Drosophila,* and unicellular ciliates, paralogous A-subunits, KATNAL1 and KATNAL2 are present, and an additional WD40-less KATNB1 paralog, KATNBL1 is present in vertebrates. KATNAL1 is highly similar in sequence and domain architecture to KATNA1^2^ and has been shown to possess MT-severing activity^4^. KATNAL2 however, outside of the conserved AAA ATPase MT severing domain, differs in domain architecture to KATNA1 and KATNAL1^2^ and in standard MT severing assays did not exhibit MT severing activity^4,5^. This aside, loss of KATNAL2 function in mouse spermatogenesis resulted in phenotypes consistent with loss of MT severing^5^.

Across eukaryotes, the regulation of cell division is a defining feature of katanin action. In *C. elegans* oocytes, the orthologues of KATNA1 and KATNB1, MEI-1 and MEI-2 drive formation of the bipolar spindle in meiosis I via MT amplification, and subsequently regulate spindle length, function and positioning^2,6^ and in meiosis II, mediate spindle disassembly^7^. KATNA1 has also been shown to regulate female meiotic spindle length in *Xenopus laevis* and *Xenopus tropocalis*^8^, and in cultured mouse oocytes KATNAL1 depletion results in abnormally long meiotic spindles and an increase in multipolar spindles ^9^. Likewise, in mitosis, katanin A-subunits have variously been shown to regulate mitotic spindle size/shape, polarity, density and orientation, and drive midbody abscission during cytokinesis in species ranging from unicellular ciliates and plants to mammalian cell lines^2,6,10^. Despite these data, the contribution of katanin proteins to male meiosis has received little attention. The requirement, and identity, of katanin A-subunits in male mammalian meiosis has not been directly tested, although data from loss-of-functions in the regulatory, B-subunit, KATNB1, strongly suggest katanin-mediated MT severing is required^11,12^.

Herein we aimed to identify the function of katanin-A subunits in male mammalian meiosis. Using a suite of single and double katanin A-subunit deletion mice, we reveal KATNA1 and KATNAL1 collectively regulate male meiosis, wherein they refine metaphase spindle structure and chromosome alignment and drive accurate chromosome segregation. We show their functions extend into haploid germ cell development (spermiogenesis), a process that necessitates the development of several specialised MT structures. We identify KATNA1 and KATNAL1 as global regulators of spermatid MTs required to restrain MT bulk, and as well as identifying several structure/process specific roles, including in Golgi function and acrosome biogenesis. We further identify KATNAL1 specific roles in cytokinesis during meiosis, in the formation of the sperm head-to-tail coupling apparatus (HTCA) and axoneme, in sperm head sharping via manchette MT dynamics and dissolution, and in sperm release (spermiation). During spermatogenesis, we reveal KATNAL1 can fully functionally compensate for loss of KATNA1. In contrast, we show KATNA1 can only partially compensate for KATNAL1 loss, and there is no functional compensation between KATNA1 and KATNAL2 in the mouse testis. Finally, using an *in vivo* immunoprecipitation (IP)-mass spectrometry (MS) approach, we reveal these diverse cellular functions are mediated by an equally diverse set of KATNA1, KATNAL1 and KATNB1 testis interacting partners, including MT associated proteins, regulators of actin dynamics and organisation, vesicle-transport proteins including multiple Rab GTPase family members, and essential mediators of male meiosis and acrosome biogenesis. Collectively these results reveal the complex reliance of spermatogenesis on katanin function and the precise neofunctionalization of A-subunits required to achieve normal sperm function and morphology.

## Results

### KATNA1 is not essential for male fertility in the mouse

Despite the clear necessity for the dynamic regulation of MT structures during meiosis, and data showing that the loss of the KATNB1 leads to notable defects in meiosis^11,12^, the ‘effector’ A-subunit is unknown. Data from *C. elegans* reveal the KATNA1 orthologue MEI-1 is a key regulator of female meiosis. To test the hypothesis that KATNA1 serves an analogous function in male meiosis, we generated a *Katna1* germ cell specific knockout mouse model (*Katna1^GCKO/GCKO^*, Fig S1A-C). As shown in Fig S1D, *Katna1* mRNA expression was reduced by 91.77% in *Katna1^GCKO/GCKO^* spermatocytes compared to *Katnal1^Flox/Flox^* controls. Loss of KATNA1 at the protein level was confirmed via immunolabelling of testis sections (Fig S1E). *Katna1^GCKO/GCKO^* mice had normal body weight (Fig S2A) and unexpectedly were fertile. *Katna1^GCKO/GCKO^* mice exhibited normal testis weight, testis daily sperm production and epididymal sperm content (Fig S2B,C,D). Histological assessment of the male reproductive tract in *Katna1^GCKO/GCKO^* mice revealed normal progression of germ cells through all phases of spermatogenesis and the presence of sperm in the epididymis (Fig S2E,F,G). Sperm from *Katna1^GCKO/GCKO^* mice were cytologically comparable to those from *Katnal1^Flox/Flox^* controls (Fig S2H) and were functionally competent, as exhibited by normal fertility when mated with wild-type females (8.2 pups per copulatory plug in *Katna1^GCKO/GCKO^* versus 7.1 pups in controls, P=0.0969). Collectively, these data reveal KATNA1 is dispensable for mammalian spermatogenesis.

### Germ cell derived KATNAL1 is required for optimal male fertility in mice

We next sought to explore the requirement in meiosis for the non-canonical katanin A-subunit, KATNAL1. A role for KATNAL1 in the somatic Sertoli cell population of the testis has been defined ^13^, but a specific role in male meiosis has not been tested. To do this we generated a *Katnal1* germ cell specific knockout mouse model (*Katnal1^GCKO/GCKO^*, Fig S3A,B). Successful ablation of *Katnal1* gene expression was confirmed by qPCR, western blotting of purified germ cells and immunolabeling of testis sections (Fig 3C-E). There was a 99.56% reduction in *Katnal1* mRNA in *Katnal1^GCKO/GCKO^* compared to *Katnal1^Flox/Flox^* germ cells.

**Figure 1:**
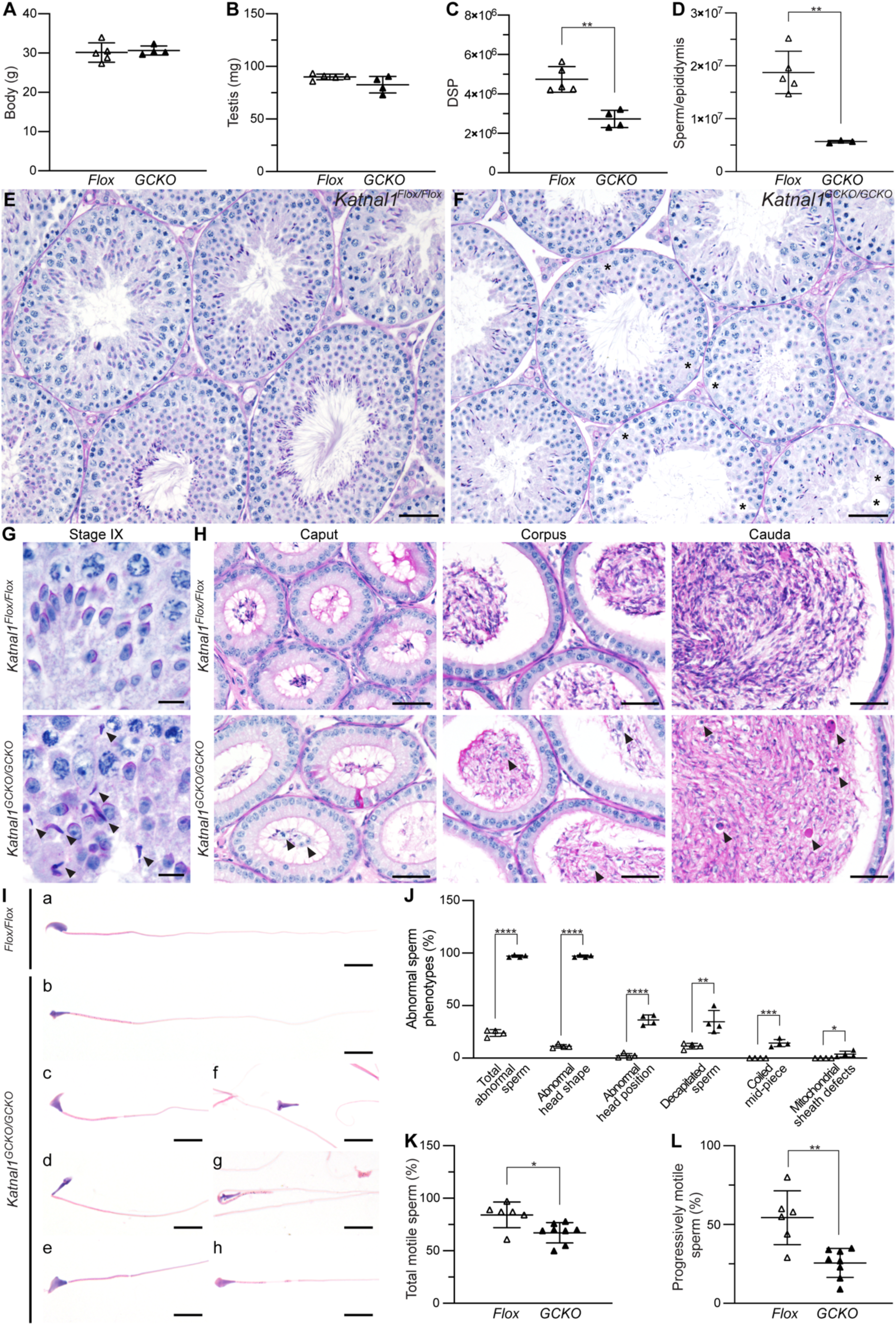
Germ cell derived KATNAL1 is required for optimal male fertility. Body weight (**A**), testis weight (**B**), DSP (**C**), and epididymal sperm content (**D**) in *Katnal1^GCKO/GCKO^* mice (black triangles) and *Katnal1^Flox/Flox^* controls (white triangles) (n*≥*3/genotype). Lines represent mean ± SD. PAS-stained testis (**E-G**) and epididymis (**H**) sections. Asterisks indicate areas of seminiferous epithelium with vacuoles or devoid of germ cells. Black arrowheads indicate retained spermatids. (**I)** Hematoxylin and eosin-stained cauda epididymal sperm. Abnormalities were observed in the vast majority of sperm from *Katnal1^GCKO/GCKO^* mice. Defects included abnormal sperm head shapes - “knobby” (b) and “hammer” head (c, f), abnormal head sperm head position (d), decapitated sperm (e,), coiled mid-pieces (g) and occasionally, mitochondrial sheath defects (h). (**J**) Quantification of abnormal cauda epididymal sperm phenotypes in *Katnal1^Flox/Flox^* (white triangles) versus *Katnal1^GCKO/GCKO^* (black triangles). (n*≥*3/genotype). Percentage of motile sperm (**K**) and percentage of progressively motile sperm (**L**) from *Katnal1^Flox/Flox^* (white triangles) versus *Katnal1^GCKO/GCKO^* (black triangles) males. Scale bars in **E-F** = 50 μm**, G** = 10 μm, **H** = 40 μm, **I** = 10 μm. \**P<*0.05 *, **P<*0.01, \*\*\**P*<0.01, \*\*\*\**P<*0.001.

**Figure 2:**
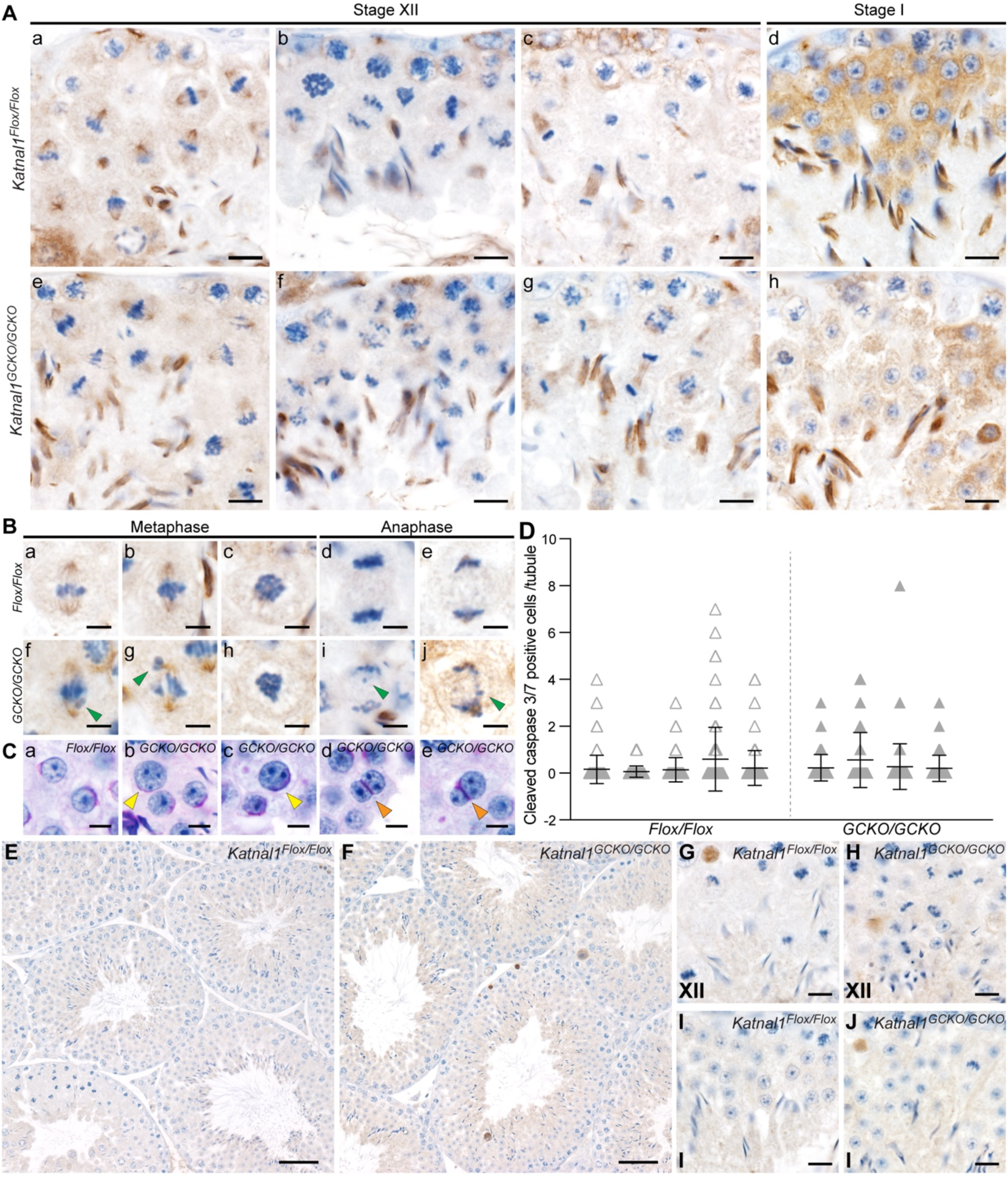
KATNAL1 regulates chromosome alignment, segregation and cytokinesis during male meiosis. (**A**-**B**) Seminiferous tubule sections immunolabelled for α-tubulin as a component of MTs. Nuclei are counterstained with hematoxylin. Green arrowheads indicate misaligned/lagging chromosomes. (**C**) PAS-stained testis sections showing abnormally large round spermatid nuclei (b,c; yellow arrowhead) and binucleated round spermatids (d-e; orange arrowheads) in *Katnal1^GCKO/GCKO^* mice. (**D**-**J**) Germ cell apoptosis in *Katnal1^GCKO/GCKO^* mice was assessed by immunolabelling of seminiferous tubules for cleaved-caspase 3 and 7. The average number of cleaved-caspase 3 and/or 7 positive germ cells per seminiferous tubule is graphed in **D** and representative images are shown in **E-G**. In **D** lines represent mean ±SD. A minimum of 80 randomly selected tubules per mouse were counted (n*≥*3/genotype). Scale bars in **A**-**B**,**G**-**J** = 10 μm**, C**=5 μm**. E**-**F**= 50 μm.

**Figure 3:**
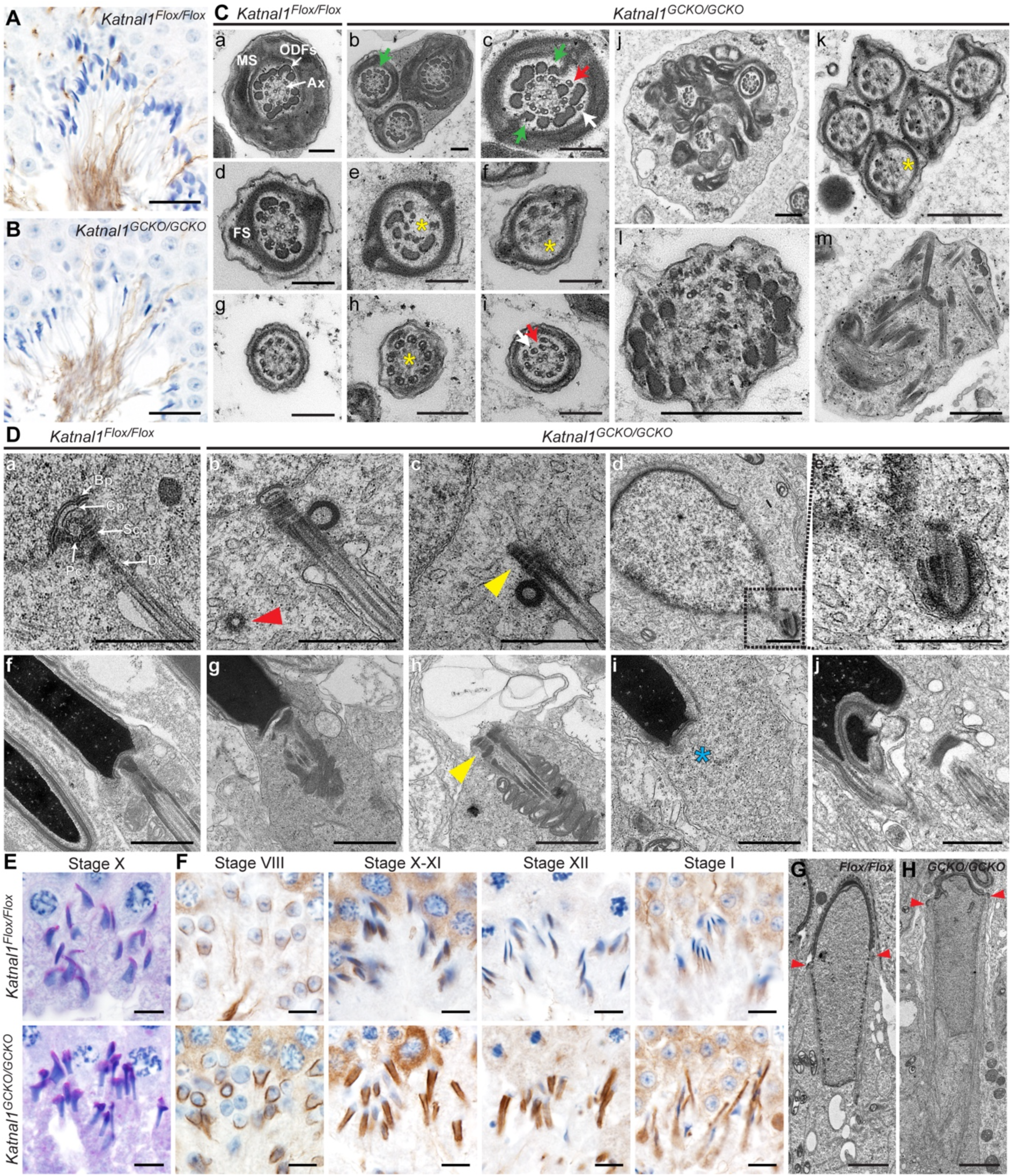
KATNAL1 is a regulator of flagella formation and the MT manchette in male germ cells. Testis sections immunolabelled for the axoneme marker acetylated tubulin in *Katnal1^Flox/Flox^* (**A**) and *Katnal1^GCKO/GCKO^* (**B**). (**C**) TEM of flagella ultrastructure in cauda epididymal sperm of *Katnal1^Flox/Flox^* and *Katnal1^GCKO/GCKO^* mice. MS= mitochondrial sheath, ODFs = outer dense fibers, Ax=axoneme FS = fibrous sheath. White arrows indicate supernumerary MT doublets. Red arrows indicate ectopic MT doublets. Green arrows indicate ectopic ODFs. Yellow asterisks indicate missing axoneme MT doublets/or central pair. (**D**) TEM of HTCA ultrastructure in round (**a**-**e**) and elongated spermatids (**f**-**j**) in *Katnal1^Flox/Flox^* and *Katnal1^GCKO/GCKO^* mice. Red arrowhead indicated supernumerary centriole. Blue asterisks indicate absent HTCA. Yellow arrowheads indicate HTCAs are abnormally detached from the nucleus. (**E**) PAS-stained testis sections showing *Katnal1^Flox/Flox^* and *Katnal1^GCKO/GCKO^* elongating spermatids. (**F**) Testis sections immunolabelled for α-tubulin as a maker of the manchette in *Katnal1^Flox/Flox^* and *Katnal1^GCKO/GCKO^* mice. Progressive steps of manchette formation, migration and dissolution are shown left to right. **(G-H)** TEM showing manchette structure in *Katnal1^Flox/Flox^* and *Katnal1^GCKO/GCKO^* elongating spermatids. Red arrowheads indicate the perinuclear ring. Scale bars in **A**-**B** = 20 μm**, Ca**-**i =** 200 nm, **Cj**-**m** = 500 nm, **D** = 1 μm, **E**-**F**=10 μm, **G**-**H** = 2 μm.

*Katnal1^GCKO/GCKO^* males exhibited normal mating frequency but were subfertile siring 84% fewer pups per copulatory plug than *Katnal1^Flox/Flox^* controls (6.9 ± 3.1SD pups / plug sired by *Katnal1^Flox/Flox^* males versus 1.4 ± 1.6SD pups / plug sired by *Katnal1^GCKO/GCKO^* males, P=0.0151). As outlined below, this subfertility was due to a dramatic reduction in sperm number, and abnormal sperm morphology and motility. *Katnal1^GCKO/GCKO^* mice had body and testis weights comparable to *Katnal1^Flox/Flox^* mice (Fig 1A,B). Likewise, the number of apoptotic germ cells per seminiferous tubules was not significantly different between *Katnal1^Flox/Flox^* and *Katnal1^GCKO/GCKO^* mice (Fig 1D). Despite this, *Katnal1^GCKO/GCKO^* testis daily sperm production was reduced by 42.3% compared to *Katnal1^Flox/Flox^* mice (Fig 1C), indicative of germ cell loss in the later steps of spermiogenesis, when spermatids contribute minimally to the overall weight of the testis. Finally, epididymal sperm content was reduced by 73.7% compared to *Katnal1^Flox/Flox^* mice (Fig 1D). This reveals that germ cell derived KATNAL1 is required for optimal spermiogenesis and, as evidenced by the more severe reduction in epididymal sperm content relative to DSPs, for their ultimate release from the seminiferous epithelium via the process of spermiation. Assessment of *Katnal1^GCKO/GCKO^* seminiferous tubule histology confirmed the abnormal retention of elongated spermatids in stage IX tubules (Fig 1G).

While all germ cell types were present in the *Katnal1^GCKO/GCKO^* seminiferous epithelium, many *Katnal1^GCKO/GCKO^* tubules contained fewer germ cells, notably spermatids, compared to *Katnal1^Flox/Flox^* mice (Fig 1E,F). Areas of Sertoli cell-only tubules were observed and gaps between adjacent Sertoli cells were frequently seen in *Katnal1^GCKO/GCKO^* but never in *Katnal1^Flox/Flox^* (Fig 1E,F). Epididymal histology was consistent with the presence of fewer sperm in *Katnal1^GCKO/GCKO^*, in addition to the presence of prematurely sloughed germ cells in *Katnal1^GCKO/GCKO^,* but not *Katnal1^Flox/Flox^*, mice (Fig 1H). Of note, immunolabelling testis sections for the Sertoli cells specific β-tubulin isoform, TUBB3, confirmed the origins of the reduced spermatogenic output in the *Katnal1^GCKO/GCKO^* were distinct from those of the previously characterised *Katnal1^1H/1H^* hypomorphic mouse model where extensive disruption of the Sertoli cell MT cytoskeleton was seen^13^. In contrast, in *Katnal1^GCKO/GCKO^* testis sections TUBB3 localisation was comparable to *Katnal1^Flox/Flox^* (Fig S4A-B). These data reveal that KATNAL1 plays an essential role in the regulation of MTs in both Sertoli and germ cells.

Of the sperm present in the epididymides of *Katnal1^GCKO/GCKO^* mice, almost all were morphologically abnormal (97.1% in *Katnal1^GCKO/GCKO^* versus 23.9% in *Katnal1^Flox/Flox^*, Fig 1I,J). Abnormalities included in head shape (97.1% in *Katnal1^GCKO/GCKO^* versus 11.3% in *Katnal1^Flox/Flox^*), head position (36.4% versus 2.3%), coupling between the head and tail (decapitation, 34.7% versus 11.5%), mid-pieces structure (coiling, 14.4% versus 0.0%) and mitochondrial sheath structure (3.9% versus 0.0%) (Fig 1J). Moreover, of those sperm found in *Katnal1^GCKO/GCKO^* epididymides, there was a 20.2% reduction in the ability for motility and a 52.9% reduction in progressive motility in the absence versus presence of KATNAL1 (Fig 2K,L). Considering the reduction in sperm number in *Katnal1^GCKO/GCKO^* mice, this equates to an 87.6% reduction in the total number of progressively motile sperm in *Katnal1^GCKO/GCKO^* compared to *Katnal1^Flox/Flox^* mice. Collectively these phenotypes reveal germ cell derived KATNAL1 is required for normal spermatogenesis and optimal male fertility.

### KATNAL1 regulates chromosome alignment, segregation and cytokinesis during male meiosis

The reduction in post-meiotic germ cells in *Katnal1^GCKO/GCKO^* seminiferous tubules indicated a variably penetrant meiotic defect. To assess *Katnal1^GCKO/GCKO^* meiotic spindles, testis sections were stained for *α*-tubulin as a component of MTs (Fig 2A,B). As seen in *Katnal1^Flox/Flox^* stage XII tubules, metaphase I and II meiotic spindle MTs typically project from the two opposing spindle poles in a narrow and uniform angle to tightly align chromosomes/chromatids along the metaphase plate (Fig 2Aa,2Ba-c). In *Katnal1^GCKO/GCKO^* stage XII tubules, the vast majority of metaphase spindles displayed normal spindle architecture and normal MT spindle density (Fig 2Ae, Bf-h). Chromosomal alignment at the metaphase plate, however, was noticeably abnormal in many *Katnal1^GCKO/GCKO^* metaphase spermatocytes, with misaligned chromosomes often observed (Fig 2A,Bf-g, green arrowheads). These defects did not however prevent progression to anaphase, with no increase in metaphase arrest and apoptosis in stage XII-I seminiferous tubules in *Katnal1^GCKO/GCKO^* compared to *Katnal1^Flox/Flox^* mice (Fig 2A,E-J).

Similarly, KATNAL1 loss in germ cells did not prevent spermatocyte progression through anaphase and telophase. Specifically there was no discernible increase in *Katnal1^GCKO/GCKO^* spermatocytes stalling in anaphase or telophase or undergoing apoptosis (Fig 2A,E-J). Instead, anaphase and telophase defects in *Katnal1^GCKO/GCKO^* meiosis led to the production of abnormal haploid germ cells. In anaphase I and II, spindle MTs undergo poleward shortening, allowing them to pull chromosomes toward opposing poles in a coordinated manner, as observed in *Katnal1^Flox/Flox^* anaphase spermatocytes (Fig 2Ab,Bd-e). In *Katnal1^GCKO/GCKO^* anaphase spermatocytes however, uneven segregation of chromosomes was often observed, with lagging chromosomes commonly seen (Fig 2Af, Bi-j, green arrowheads). Consistent with uneven chromosome segregation in anaphase, many round spermatids possessed abnormally large nuclei (Fig 2Cb-c, yellow arrowheads). During telophase and cytokinesis midbodies were present in *Katnal1^GCKO/GCKO^* and were comparable to those in *Katnal1^Flox/Flox^* spermatocytes (Fig 2Ac,g). However, a subset of the resulting spermatids was binucleated (Fig 2Cd,e, orange arrowheads), and as detailed below sperm/spermatids with multiple axonemes (Fig 3C) and supernumerary centrioles (Fig 3Db, red arrowhead) were often observed. This indicates that while midbodies do form in the absence of *Katnal1*, KATNAL1 plays a key role its remodelling into an intracytoplasmic bridge and the maintenance of individual daughter spermatids. Binucleated spermatids were never observed in *Katnal1^Flox/Flox^* tubules.

These data reveal KATNAL1 regulates MT-chromosome attachment and/or metaphase spindle dynamics but is dispensable for MT generation and pruning during spindle biogenesis in male meiosis. Moreover, our data show KATNAL1 greatly facilitates accurate chromosome segregation during anaphase and midbody function and disassembly during cytokinesis in either meiosis I and/or meiosis II.

### KATNAL1 regulates flagella and manchette formation in male germ cells

As detailed above, of the epididymal sperm that were produced in *Katnal1^GCKO/GCKO^* mice 97.1% were morphologically abnormal revealing that KATNAL1 is required during spermatid differentiation (spermiogenesis) (Fig 1I,J). One of the first processes to commence in spermiogenesis is acrosome formation, and our recent data identified MT severing as a potential regulator of this process ^11^. The identity of the involved MT severing A-subunit remains unknown. Acrosome formation was overtly normal in *Katnal1^GCKO/GCKO^* mice compared to *Katnal1^Flox/Flox^* controls, indicating KATNAL1 is not essential for this process (Fig S4C).

Multiple abnormalities were however observed in *Katnal1^GCKO/GCKO^* mouse sperm tails and the head-to-tail coupling apparatus (HTCA), including abnormal reticulation of the sperm head, decapitation, coiling of the sperm tail mid-piece and mitochondrial sheath defects, in addition to reduced sperm motility. Staining of *Katnal1^GCKO/GCKO^* testis sections for acetylated tubulin, as a marker of stable MTs, revealed axonemal MTs were present in sperm tails (Fig 3B), however analysis via transmission electron microscopy (TEM) revealed a range of axoneme and accessory structure abnormalities (Fig 3C). In *Katnal1^GCKO/GCKO^* mice, sperm flagella axonemes often lacked the typical 9+2 MT doublet arrangement observed in controls (Fig 3Ca,d,g). Instead MT doublets were often missing (Fig 3Ce,f,h, yellow asterisks), ectopically located (Fig 3C, red arrowheads) or in many cases supernumerary MT doublets were present (Fig 3Cc,l, white arrowheads). In the most dramatic instances, and consistent with the failures in cytokinesis, multiple axonemes, often each with associated accessory structures, were observed within one cell (Fig 3Cb,j-l). The abnormalities in MT doublet number and location were typically accompanied by defects in outer dense fiber (ODF) number and location (Fig 3C). In wild-type sperm, the mid-piece contains 9 ODFs closely associated with the 9 outer MT doublets of the axoneme (Fig 3Ca). In the principal piece ODFs 3 and 8 are replaced by the longitudinal columns of the fibrous sheath (Fig 3Cd). In *Katnal1^GCKO/GCKO^* mice however, sperm tail ODFs 3 and/or 9 frequently persisted into the principal piece (Fig 3Cb,c, green arrowheads). In *Katnal1^GCKO/GCKO^* mouse sperm tails wherein some or all axoneme MTs were missing, ODFs were often also missing (Fig 3Ce,f). Consistent with the coiled mid-pieces and mitochondrial sheath defects observed in some sperm from *Katnal1^GCKO/GCKO^* mice, TEM revealed that in many sperm the mitochondria were not properly organised into a helical sheath (Fig 3Cj). Fibrous sheath development in *Katnal1^GCKO/GCKO^* sperm however, appeared normal in most cells (Fig 3C). Finally, in some sperm, only disorganised fragments of sperm tail components were observed indicative of degenerating cells (Fig 3Cm). Collectively, these data reveal KATNAL1-mediated MT severing regulates the assembly and potentially stability of axoneme MT doublets, and that KATNAL1 contributes to the development of the mitochondrial sheath and ODFs, but not the fibrous sheath. These data are consistent with defects in cargo/organelle delivery to and along the growing sperm tail. Indeed, there is an increasing of body of evidence (reviewed by Pleuger et al.^14^) that mitochondria and ODFs rely on MT-based transport mechanisms for their delivery into the sperm tail.

The frequent decapitation or hyper-extension of sperm heads, in sperm from *Katnal1^GCKO/GCKO^* males, indicates the HTCA was weak and/or improperly formed in the absence of KATNAL1. Moreover, TEM of round and elongated spermatids revealed a range of HTCA abnormalities in *Katnal1^GCKO/GCKO^* mice compared to controls (Fig 3D). In *Katnal1^Flox/Flox^* mice, spermatid HTCAs were easy to find in testis sections via TEM. As expected, they contained a proximal centriole coupled to the nucleus via the basal plate and capitulum, a distal centriole coupled to the plasma membrane and continuous with the axoneme, and the surrounding segmented columns which are continuous with the ODFs (Fig 3Da,f). In *Katnal1^GCKO/GCKO^* mice, however normal HTCAs were rare (Fig 3Db,g). In most spermatids, HTCAs were abnormal (Fig 3Dc-e, h-j). In some cells, the HTCA and associated axoneme was present, but not attached to the nucleus (Fig 3Dc,h, yellow arrowheads), while in others the basal plate could be seen associated with the nucleus, but the rest of the HTCA was absent (Fig 3Dg). Consistent with this data, immunolabelling of sperm for SUN5, which is an essential component of the LINC complex that physically docks the sperm centrosome to the nucleus, revealed a 57.2% reduction in sperm from *Katnal1^GCKO/GCKO^* males containing SUN5 at the HTCA compared to sperm from *Katnal1^Flox/Flox^* mice (Fig S4D,E). These data suggest that KATNAL1 regulated-MTs are either required to fortify the connection between the basal plate and the nucleus or are required for the delivery and deposition of essential components of the HTCA. Our data also suggest that SUN5 is delivered to the HTCA via KATNAL1-regulated MTs.

Abnormal head shape was the most frequent morphological abnormality present in the sperm of *Katnal1^GCKO/GCKO^* mice. Instead of the normal hook shape exhibited in 88.75% of *Katnal1^Flox/Flox^* mouse sperm, 34% of *Katnal1^GCKO/GCKO^* mouse sperm heads exhibited a ‘hammerhead’ shape and 62% exhibited a ‘knobby’ head phenotype (Fig 1I, S4F,G). Abnormal head shaping became apparent during nuclear elongation (Fig 3E) and were consistent with defects in manchette function, where the manchette is a transient MT-based structure which sculpts the distal half of the spermatid head. As shown in Fig 3F in wild-type mice the manchette forms in step 8-9 spermatids (stage VIII-IX seminiferous tubules), before migrating distally while simultaneously constricting to shape the spermatids head during steps 10-12 (stages X-XII; Fig 3F). In steps 13-14 (stage I-III) the manchette is dissembled (Fig 4F). In spermatids from *Katnal1^GCKO/GCKO^* mice, manchettes formed at step 8-9 but were abnormally elongated. Elongation became increasingly pronounced as spermatid development continued and was associated with a failure of manchette perinuclear ring migration down the sperm head, but continued constriction, and resulting in a ‘knobby head’ phenotype (Fig 4G). Finally, manchette disassembly in steps 13-14 was delayed in the absence of KATNAL1 (Fig 4F). Collectively, these results reveal KATNAL1-mediated MT severing is required to regulate manchette MT length, migration and disassembly.

**Figure 4:**
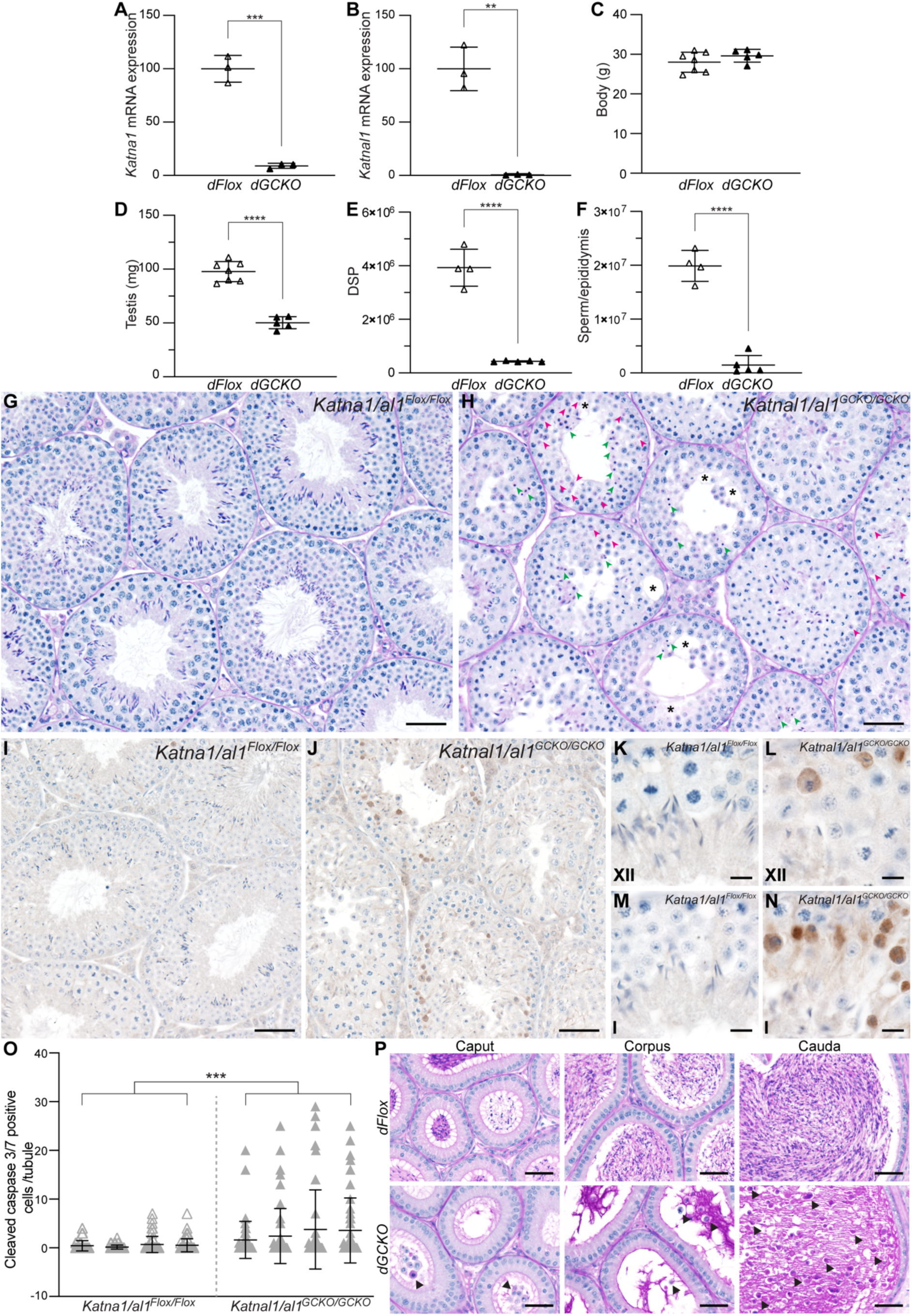
Double germ cell deletion of KATNA1 and KATNAL1 results in a severe oligozoospermia-like phenotype. qPCR analysis of *Katna1* (**A**) and *Katnal1* (**B**) transcript levels in *Katna1/al1^GCKO/GCKO^* (black triangles, *dGCKO*) isolated spermatocytes relative to *Katna1*/*al1^Flox/Flox^* (white triangles, *dFlox*) isolated spermatocytes (n*≥*3/genotype). Body weight (**C**), testis weight (**D**), testis DSP (**E**), and epididymal sperm content (**F**) in *Katna1/al1^GCKO/GCKO^* (black triangles) compared to *Katna1/al1^Flox/Flox^* controls (white triangles) (n*≥*3/genotype). Lines represent mean ± SD. PAS-stained testis (**G-H**) sections from *Katna1/al1^GCKO/GCKO^* and *Katna1/al1^Flox/Flox^* mice. Magenta arrowheads indicate pyknotic spermatocytes and green arrowheads indicate pyknotic spermatids. Asterisks indicate areas of seminiferous epithelium with vacuoles or devoid of germ cells. Analysis of germ cell apoptosis in *Katnal1^GCKO/GCKO^* mice relative to *Katna1/al1^Flox/Flox^* mice (**I**-**O**). Representative images of testis immunolabelling of cleaved-caspase 3 and 7 are shown in (**I**-**N**). Roman numerals denote seminiferous tubule stage. The average number of cleaved-caspase 3 and/or 7 positive germ cells per seminiferous tubule per mouse is graphed in (**O**). In **O** lines represent mean ±SD. A minimum of 50 randomly selected tubules per mouse were counted (n*≥*3/genotype). PAS-stained epididymis sections (**P**) from *Katna1/al1^GCKO/GCKO^* and *Katna1/al1^Flox/Flox^* mice. Scale bars in **G**-**J** = 50 μm, in **K**-**N** = 10 μm, **P =** 40 μm. \**P<*0.05 *, **P<*0.01, \*\*\**P*<0.01, \*\*\*\**P<*0.001.

### KATNA1 and KATNAL1 compensate for each other in male meiosis and spermiogenesis

Given the apparently normal fertility in *Katna1^GCKO/GCKO^* and residual fertility in *Katnal1^GCKO/GCKO^* mice, we sought to explore the possibility of functional compensation, between KATNA1 and its paralogs KATNAL1 and KATNAL2 in the testis. This is a particularly critical question for meiosis wherein the identity of the critical A-subunit(s) remains unknown. As reported previously, *Katnal2* loss has no apparent effect on male meiosis but does result in male infertility due to severe sperm developmental defects^5^. To test for redundancy, we generated (1) *Katna1* and *Katnal1* double germ cell knockout mice (*Katna1/al1^GCKO/GCKO^*) and (2) double *Katna1* germ cell and *Katnal2* knockout mice (*Katna1^GCKO/GCKO^/al2^KO/KO^*). The *Katna1^GCKO/GCKO^/al2^KO/KO^* mouse model (Fig S5) phenocopied our previously characterised *Katnal2* mutant models^5^, including having a comparable spermiation failure phenotype, sperm head shaping defects (knobby heads), and an absence of normal sperm tail development (Fig S5). No aspect of the infertility phenotype appeared worse that the loss of KATNAL2 in isolation. These data reveal that KATNA1 and KATNAL2 do not have compensatory function in male germ cell development.

By contrast the removal of KATNA1 and KATNAL1 from germ cells resulted in a significantly worse male fertility phenotype than seen in either individual knockout, especially in meiosis. Successful ablation of both *Katna1* and *Katnal1* from male germ cells was confirmed by qPCR (Fig 4A,B). Test mating of the *Katna1/al1^GCKO/GCKO^* male mice with wild-type females, revealed loss of both KATNA1 and KATNAL1 from male germ cells results in male sterility (7.1±3.2 SD pups / plug sired by *Katna1/al1^Flox/Flox^* males versus 0± 0 SD pups / plug sired by *Katna1/al1^GCKO/GCKO^* males (P=0.0175)). Moreover, assessment of the reproductive tract revealed a 48.5% reduction in testis weight, an 89% reduction in daily sperm output and a 96.2% reduction in epididymal sperm content compared to controls (Fig 4D-F). Examination of *Katna1/al1^GCKO/GCKO^* seminiferous tubule histology instantly revealed large scale germ cell loss during both meiosis and spermiogenesis that was significantly worse than in either single gene deletion (Fig 4G-H). Numerous pyknotic spermatocytes were observed in all stage XII and I *Katna1/al1^GCKO/GCKO^* tubules indicative of cells stalling in meiosis followed by apoptosis but were rare in *Katna1/al1^Flox/Flox^* (Fig 4G-H, magenta arrowheads). Of the germ cells that progressed through meiosis in *Katna1/al1^GCKO/GCKO^*, the vast majority became pyknotic by steps 12-13 (stages XII-I) of elongating spermatid development (Fig 4H, green arrowheads) and as a result elongated spermatids were absent past stage II-VIII *Katna1/al1^GCKO/GCKO^* tubules (Fig 4H). Moreover, analysis of cleaved caspase 3/7 immunolabelling revealed a 6.4 fold increase in the number of apoptotic germ cells in *Katna1/al1^GCKO/GCKO^* versus *Katna1/al1^Flox/Flox^* seminiferous tubules (Fig 4O). The majority of apoptotic cells were spermatocytes that had stalled in anaphase I and meiosis II in stage I tubules (Fig 5J,N), and to a lesser degree those undergoing metaphase I in stage XII (Fig 5J,N). In line with these defects, analysis of epididymal sections confirmed almost no sperm reached the *Katna1/al1^GCKO/GCKO^* epididymis. The bulk of the cells in the lumen of *Katna1/al1^GCKO/GCKO^* epididymides were instead prematurely sloughed round germ cells (Fig 4P, arrowheads).

**Figure 5:**
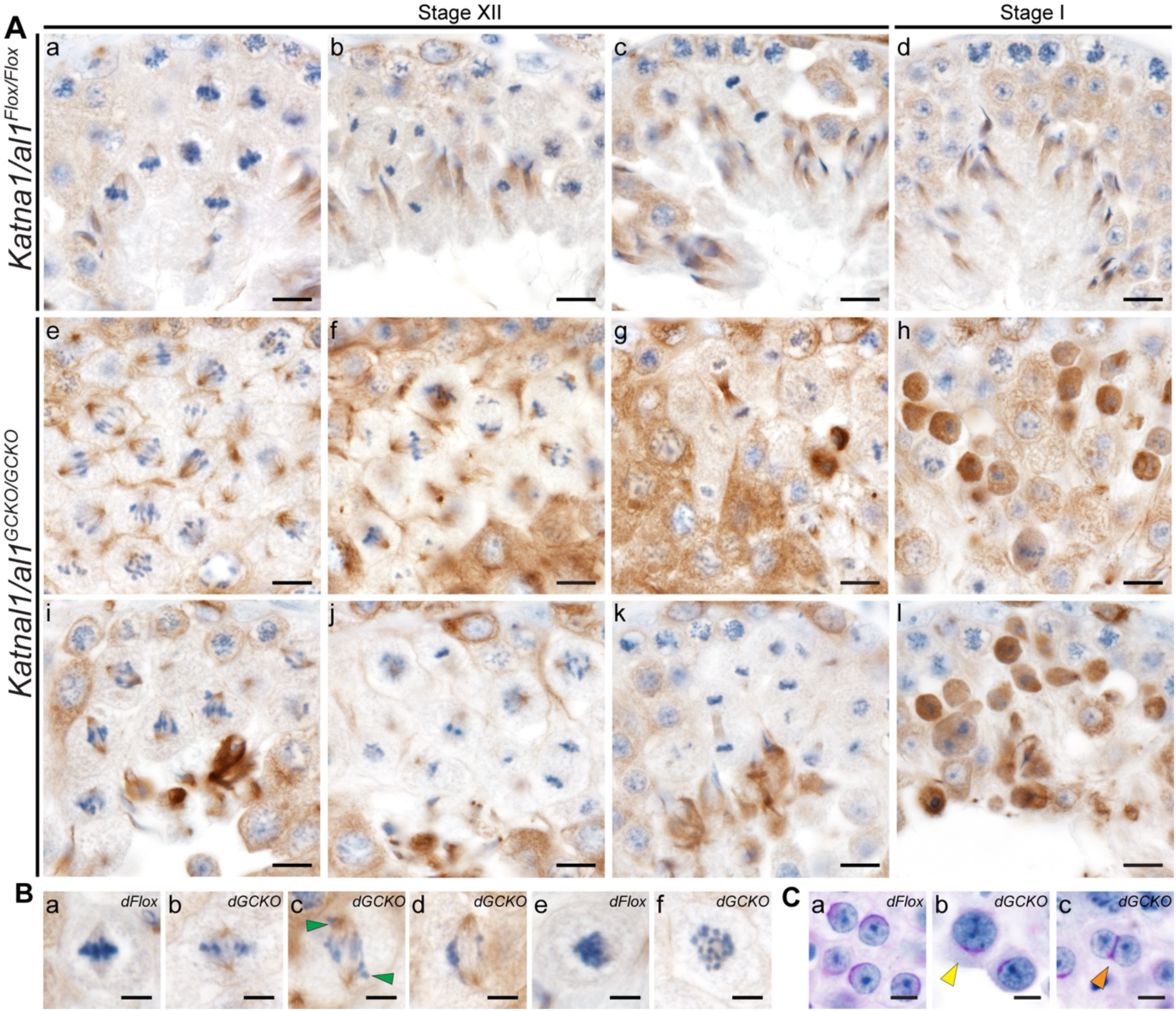
KATNA1 and KATNAL1 compensate for each other in male meiosis. (**A**-**B**) *Katnal1/al1^Flox/Flox^* (*dFlox*) and *Katna1/al1^GCKO/GCKO^* (*dGCKO*) seminiferous tubule sections immunolabelled for MTs (α-tubulin). Nuclei are counterstained with hematoxylin. Green arrowheads indicate misaligned/lagging chromosomes. (**C**) PAS-stained *Katnal1/al1^Flox/Flox^* (*dFlox*) and *Katna1/al1^GCKO/GCKO^* (*dGCKO*) testis sections showing abnormally large round spermatid nuclei (b; yellow arrowhead) and binucleated round spermatids (c; orange arrowheads) in *Katna1/al1^GCKO/GCKO^* mice. Scale bars in **A**= 10 μm, in **B**-**C**= 5 μm.

Assessment of *Katna1/al1^GCKO/GCKO^* meiotic spindles, via the immunolabelling of testis sections for *α*-tubulin, revealed that while meiotic spindles formed in *Katna1/al1^GCKO/GCKO^* spermatocytes, most were morphologically abnormal compared to *Katna1/al1^Flox/Flox^* controls (Fig 5A,B). At metaphase, in contrast to the tightly packed morphology of *Katna1/al1^Flox/Flox^* controls (Fig 5Aa,Ba,e), chromosomes were loosely dispersed along the metaphase plate in *Katna1/al1^GCKO/GCKO^* mice (4Bb,f). Misaligned and/or lagging chromosomes were frequently observed in *Katna1/al1^GCKO/GCKO^* metaphase and anaphase spermatocytes (Fig 5A,B). As detailed above, these defects ultimately lead to a large proportion of *Katna1/al1^GCKO/GCKO^* metaphase and anaphase spermatocytes stalling and undergoing apoptosis in stage XII-I seminiferous tubules (Fig 4H, I-O). Of note, examination of *Katna1/al1^GCKO/GCKO^* stage I tubules, revealed that these stalled metaphase and anaphase spermatocytes where characterised by the ectopic accumulation of *α*-tubulin throughout their cytoplasm (Fig 5Ah,i) indicating that KATNA1/KATNAL1-mediated severing restrains MT bulk in spermatocytes. Of the subset of *Katna1/al1^GCKO/GCKO^* spermatocytes that progressed through meiosis, overtly normal midbody formation was observed in telophase (Fig 5Ag,k). However, indicative of failure of midbody to intracellular bridge conversion and similar to *Katna1^GCKO/GCKO^* mice, binucleated spermatids were frequently observed in *Katna1/al1^GCKO/GCKO^* mice but never in *Katna1/al1^Flox/Flox^* controls (Fig 5Ag,k). Consistent with compromised anaphase chromosome segregation, abnormally large round spermatids were frequent in *Katna1/al1^GCKO/GCKO^* mice but never in *Katna1/al1^Flox/Flox^* controls (Fig 5Ag,h). Collectively these data indicate that KATNAL1 and KATNA1 can at least partially compensate for each other in metaphase and anaphase of male meiosis. The frequency of binucleated spermatids in *Katna1/al1^GCKO/GCKO^* mice however, was similar to that of *Katnal1^GCKO/GCKO^* suggesting KATNA1 does not compensate for KATNAL1 in cytokinesis.

Assessment of spermiogenesis in *Katna1/al1^GCKO/GCKO^* mice revealed that unlike the single GCKO mice, acrosome biogenesis was overtly abnormal in the absence of both KATNA1 and KATNAL1 indicating functional compensation between these two A-subunits (Fig 6A,B). In PAS-stained *Katna1/al1^Flox/Flox^* control testis sections, a single PAS-positive acrosomal vesicle was observed to adhere to step 2-3 round spermatid nuclei (stage II-III) in the Golgi phase of acrosome development, and progressively spread to cover the apical half of the spermatid nuclei as spermiogenesis progressed (Fig 6A). In *Katna1/al1^GCKO/GCKO^* mice however, a large subset of round spermatids with supernumerary ectopic acrosomal vesicles were observed (Fig 6A orange arrowheads), in addition to a smaller subset of round spermatids wherein pro-acrosomal vesicles had failed to attach to the nucleus at the appropriate time (Fig 6A, yellow arrowheads). As spermiogenesis progressed, elongating spermatids with fragmented and partially detached acrosomes were common in *Katna1/al1^GCKO/GCKO^*, but never seen in *Katna1/al1^Flox/Flox^* controls (Fig 6A, blue arrowheads). Ectopic PAS-positive vesicles were also present throughout the seminiferous epithelium during multiple steps of acrosome biogenesis in *Katna1/al1^GCKO/GCKO^*, but not *Katna1/al1^Flox/Flox^*, mice (Fig 6A, green arrowheads). TEM of acrosome biogenesis confirmed these observations (Fig 6B), and revealed morphological abnormalities in the spermatid Golgi apparatus of *Katna1/al1^GCKO/GCKO^* mice (Fig 6Bi,k) compared to *Katna1/al1^Flox/Flox^* mice (Fig 6Bc,e). In normal round spermatids, and as seen in *Katna1/al1^Flox/Flox^* mice (Fig 6Bc,e), the Golgi apparatus has a flattened sometimes horseshoe-like morphology wherein the trans-Golgi network (TGN) is closely associated with, and oriented in parallel to, the apical nuclear membrane. In *Katna1/al1^GCKO/GCKO^* mice however the Golgi stacks were disorganised, often displaying a circular morphology, wherein the cis-Golgi network almost entirely enclosed the TGN (Fig 6Bi), or a dispersed, splayed morphology (Fig 6Bk). These phenotypes, reveal KATNA1 and KATNAL1 collectively regulate the spatial organisation of the Golgi apparatus and the transport of proacrosomal vesicles, and their fusion, to the nuclear membrane. The acrosome abnormalities observed in *Katna1/al1^GCKO/GCKO^* mice, are reminiscent of those seen in KATNB1 loss of function mice^11^ suggesting these KATNA1 and KATNAL1 likely function in complexes with KATNB1 during acrosome biogenesis.

**Figure 6:**
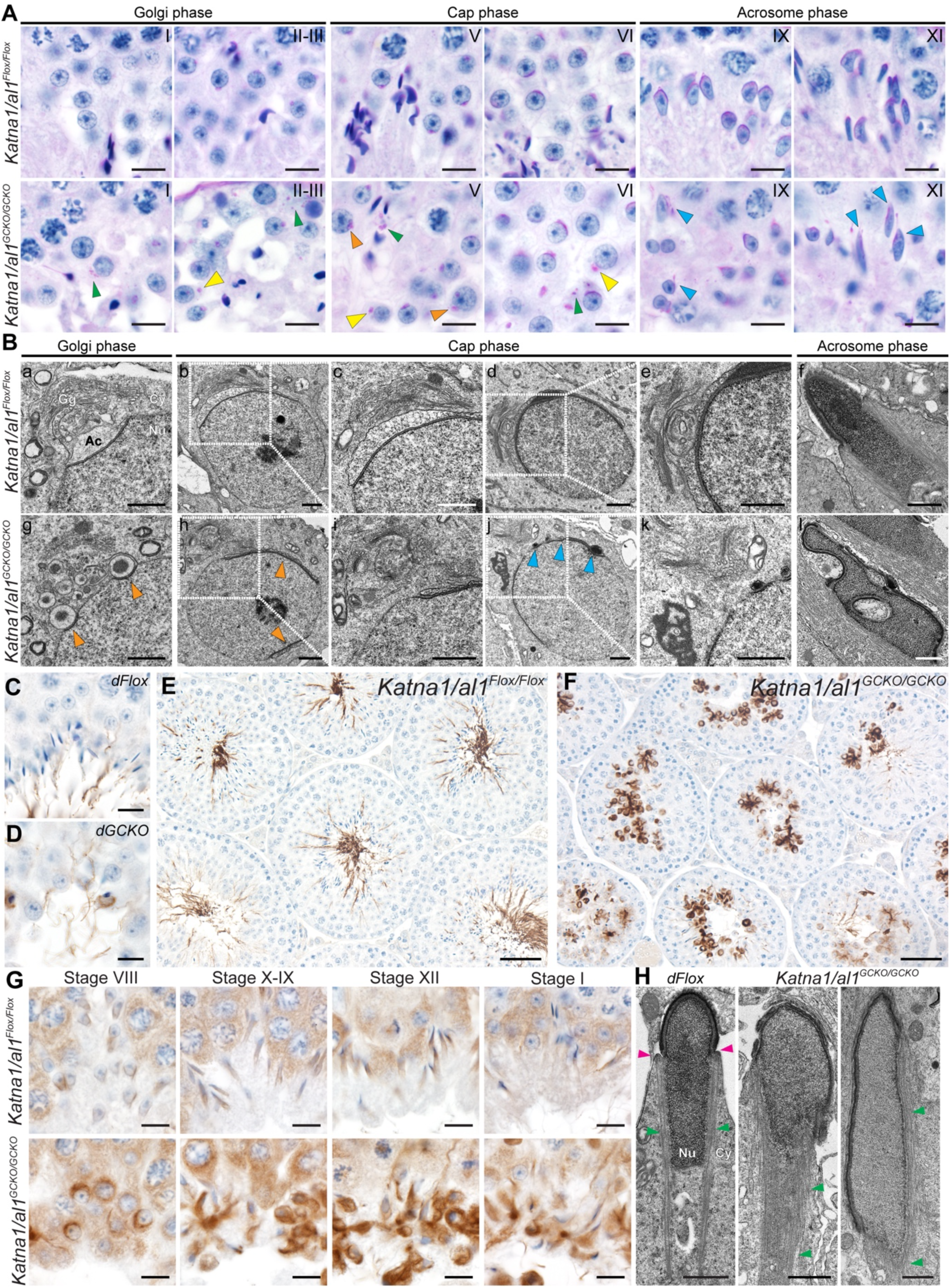
KATNA1 and KATNAL1 compensation in spermatid acrosome formation and MT regulation. (**A**) PAS-stained testis sections showing acrosome formation in *Katna1/al1^Flox/Flox^* and *Katna1/al1^GCKO/GCKO^* mice. Progressive steps of acrosome formation are shown left to right. Roman numerals denote seminiferous tubule stage. Orange arrowheads indicate nuclei with more than one proacrosomal vesicle attached. Yellow arrowheads indicate acrosomal vesicles not attached to the nucleus. Green arrowheads indicate ectopic PAS positive vesicles abnormally dispersed in round spermatid cytoplasm. Blue arrowheads indicate elongating spermatids with abnormal gaps/fragmentation of the acrosomes. (**B**) Electron micrographs showing acrosome and Golgi apparatus ultrastructure in *Katna1/al1^Flox/Flox^* and *Katna1/al1^GCKO/GCKO^* mice. Orange arrowheads indicate supernumerary site of acrosome formation. Blue arrowheads indicate numerous small unfused acrosomal vesicles docked at the nuclear membrane/fragmentation of the acrosome. (**C**-**F**) *Katna1/al1^Flox/Flox^* (*dFlox*) and *Katna1/al1^GCKO/GCKO^* (*dGCKO*) mouse testis sections immunolabelled for the sperm tails marker acetylated tubulin. (**G**) *Katna1/al1^Flox/Flox^* and *Katna1/al1^GCKO/GCKO^* testis sections immunolabelled for α-tubulin as a maker of the manchette. Progressive steps of manchette formation, migration and dissolution are shown left to right. TEM showing nuclear and manchette morphology in *Katna1/al1^Flox/Flox^* (*dFlox*) and *Katnal1^GCKO/GCKO^* mouse elongating spermatids. Red arrowheads indicate the perinuclear ring. Blue arrowheads indicate MTs. Scale bars in **A, C-D** = 10 μm, **B,H** = 1 μm, **E-F** = 50 μm, **G**=10 μm. Nu = Nucleus, Cy =cytoplasm Gg=Golgi apparatus, Ac=acrosome.

Analysis of the key MT-based structures required for spermiogenesis revealed phenotypes consistent with a loss of MT severing and global dysregulation of the MT cytoskeleton. Similar to *Katna1^GCKO/GCKO^* mice, immunolabelling of testis sections for acetylated tubulin revealed sperm tails formed in *Katna1/al1^GCKO/GCKO^* mice, however reflective of sperm numbers fewer tails were seen than in *Katna1/al1^Flox/Flox^* controls (Fig 6C-F). Acetylated tubulin staining however, revealed the frequent ectopic accumulation of stable MTs in the cytoplasm of spermatids in *Katna1/al1^GCKO/GCKO^* mice (Fig 6E-F), consistent with the decreased turnover of MTs.

Assessment of manchette formation via the immunolabelling of testis sections for *α*-tubulin revealed a catastrophic dysregulation of manchette formation. MTs accumulated around the spermatid nucleus at stage VIII in *Katna1/al1^GCKO/GCKO^* mice (Fig 6G), however they failed to assemble into a sheath around, and with MTs aligning in parallel to, the distal half of the nucleus. Instead, the cytoplasm of the elongating spermatids became enriched in ectopic *α*-tubulin as spermiogenesis progressed (Fig 6G). Analysis of elongating spermatid ultrastructure via TEM confirmed these observations (Fig 6H). In *Katna1/al1^Flox/Flox^* mouse elongating spermatids, a typical manchette structure wherein MTs projected distally from a perinuclear ring was observed (Fig 6H). In *Katna1/al1^GCKO/GCKO^* mice, MTs were observed throughout the cytoplasm of elongating spermatids, however these were never associated with a perinuclear ring and were not arranged into typical manchette structure (Fig 6H). Collectively, these results reveal that KATNA1/KATNAL1-mediated severing is required to restrain MT growth/prune MT bulk during spermatid differentiation and to regulate manchette formation. Moreover, the absence of these phenotypes in the single GCKO KATNA1 and KATNAL1 models, suggest KATNA1 and KATNAL1 can compensate for each other in these aspects of spermiogenesis.

### KATNA1, KATNAL1 and KATNB1 interact with a range of cytoskeletal and vesicular transport proteins in the mouse testis

To explore the mechanism of action of KATNA1 and KATNAL1 during male meiosis and spermiogenesis, and how they functionally compensate for each other, we determined the respective interactome for KATNA1, KATNAL1 and KATNB1, within testis immunoprecipitates via MS (Table S1 and Fig S6A-C). A total of 322, 108 and 213 proteins were significantly enriched in the KATNA1, KATNAL1 and KATNB1 IP respectively compared to their corresponding controls (Table S1 and Fig S6A-C). As expected, KATNA1, KATNAL1 and KATNB1 were all significantly enriched in their own IPs validating the specificity of each IP. KATNA1 and KATNAL1 were not found in the binding partner list for each other, suggesting that they do not interact with each other to form heterohexamers (Table S1). Similarly, neither bound to KATNAL2, however KATNAL1 and KATNB1 each pulled the other down (Table S1). Consistent with the three proteins having similar functions, they shared 48 candidate interacting proteins, in addition to 102 candidates that were shared by two katanins (Fig S6D).

PANTHER protein class and gene ontology analysis^15^ was conducted (Table S2 and summarized in Fig S6E-J). Consistent with their regulation of MT-based processes in spermatogenesis, KATNA1, KATNAL1 and KATNB1 each bound several cytoskeletal proteins (Fig S6E-G, Table S2). Of particular relevance to the regulation of the Golgi and acrosome formation all bound proteins involved in vesicle-mediated transport. In addition, KATNA1 bound to known vesicle targeting proteins including ARL3 (Fig S6E-G, Table S2). Functional descriptions of selected cytoskeletal, membrane/vesicle-transport and spermatogenesis related proteins significantly enriched in the KATNA1, KATNAL1 and KATNB1 IP experiments are summarised in Table S3.

Strikingly, across the three IP experiments the cytoskeletal proteins identified were predominantly actin-related proteins, however tubulin and MT-related proteins were also identified (Fig S6E-J, Table S2 and S3). KATNA1 also bound to the intermediate filament protein vimentin (Table S2 and S3). Of the MT-related proteins identified as katanin testis-interactors, multiple proteins involved in the regulation and activity of dynein MT motor proteins were identified. Indeed, KATNA1 and KATNB1 both bound components of the dynactin complex (Table 1), an essential cytoplasmic-dynein cofactor involved in MT-based cargo transport. KATNA1 also bound DNAAF4, which is required for axonemal dynein assembly and thus ciliary motility, and all three katanins bound NUDC, a dynein-dynactin complex regulator (Table S3). Consistent with the phenotypes of our *Katnb1*^11,12^, *Katnal1* and *Katna1/al1* mutant mice, the MT-related proteins identified included those with established functions in the mitotic and/or meiotic spindle (e.g. NUDC, MAPRE1/EB1), intraflagellar transport and ciliogenesis (ARL3) and axoneme assembly DNAAF4; Table S3). Moreover, several have previously been implicated in spermatogenesis and male fertility (e.g. ARL3, DNAAF4, MAPRE1/EB1) and, consistent with its mechanism of action, KATNA1 pulled down both *α*- and *β*-tubulin isoforms (Table S3).

Of the actin-binding proteins identified across the three katanin IP experiments, multiple regulators of actin polymerization/dynamics were identified including components of the ARP2/3 and CapZ complexes (Table S2, S3). KATNA1, KATNAL1 and KATNB1 each bound at least two components of the ARP2/3 complex. Data strongly suggests this complex plays key roles in blood-testis-barrier integrity, spermiogenesis and spermiation^16–19^. Likewise, all bound the CapZ subunit CAPZA1, and KATNA1 and KATNAL1 both bound CAPZA2. In addition, several regulators of actin-network organization were identified to bind KATNA1 and KATNB1, and KATNA1 bound MYL6, an actin motor protein which localizes to the manchette (Table S3). Collectively, these data indicate a novel connection between katanin proteins and the actin cytoskeleton in male germ cells. This is consistent with data showing the *Arabidopsis* katanin A-subunit KATANIN1 regulates the actin cytoskeleton^20^ and that cortical localisation of *Drosophila* KATNA1 orthologue kat-60 is actin-dependent^21^.

KATNA1 and KATNB1 bound multiple Rab GTPases, a family which regulates multiple aspects of vesicle transport (vesicle budding, actin/MT-mediated transport and fusion) (Tables 1, S1 and S2). Of note in spermatogenesis there is increasing evidence Rab and Rab-like proteins mediate pro-acrosomal vesicle transport, in addition to intraflagellar transport and sperm tail development^22–25^. Other transport proteins included those related to endoplasmic reticulum (ER) to Cis-Golgi transport, intra-Golgi transport, clathrin-mediated vesicle disassembly and endocytosis (Table 1). Further consistent with the acrosome defects in *Katna1/al1^GCKO/GCKO^* mice, KATNAL1 bound ACRBP a protein essential for acrosome granule formation and maintenance of acrosome integrity, and consistent the Golgi defects, a number of proteins were identified with roles in Golgi organization (e.g. Rab2, Rab6, VCP, LMAN1; Table S3).

Other notable proteins identified as binding the katanins included essential male meiosis regulators TEX15 (KATNA1 IP) and HSPA2 (KATNA1/B1 IP), and FABP9 a protein required for normal sperm head shape in mice (KATNA1/AL1 IP) (Table S3).

## Discussion

Katanin MT severing enzymes are potent regulators of MT dynamics and organization across eukaryotes. Herein, using a combination of single and double katanin A-subunit GCKO mice, we show for the first time the essential requirement for katanin A-subunits in mammalian male meiosis, provide the first direct evidence that different katanin A-subunits can functionally compensate for one another and establish KATNA1 and KATNAL1 as novel regulators of spermatid differentiation (spermiogenesis). Specifically, KATNA1 and KATNAL1 collectively regulate spermatocyte metaphase and anaphase spindle structure and function, acrosome development via the Golgi apparatus and pro-acrosomal vesicle trafficking, sperm head shaping via manchette formation, and collectively restrain MT bulk during male meiosis and spermiogenesis. In addition to this, KATNAL1 regulates spermatocyte cytokinesis, sperm tail and HTCA development, manchette dynamics and dissolution, and the release of sperm from the seminiferous epithelium (spermiation). Through the analysis of KATNA1, KATNAL1, and KATNB1 testis binding proteins, we reveal these activities involve interactions with a diverse suite of proteins, including actin-, MT-, vesicle/membrane trafficking-, and spermatogenesis-related proteins.

The finding that KATNA1 and KATNAL1, but not KATNA1 and KATNAL2, have gene redundancy in mammalian spermatogenesis is consistent with the highly similar domain structure and severing activity shared between KATNA1 and KATNAL1, but not KATNAL2^2^. Interestingly, KATNAL1 fully compensates for KATNA1 germ cell loss, whereas KATNA1 can only partially compensate for germ cell KATNAL1 loss. This suggests KATNAL1 is the more potent/efficient MT severing enzyme in the testis, and/or that it has neofunctionalized to acquire additional functions. We predict it is a combination of the two. Indeed, KATNAL1 has possesses higher MT severing activity in mouse Neuro-2a cells, and is less prone to degradation in HEK293T cells compared to KATNA1^26^. However, we also identified several different KATNA1 versus KATNAL1 testis interacting proteins indicating they have at least some distinct modes of regulation and/or function. These findings have important implications for the study of katanin proteins in contexts wherein multiple A-subunits exist, revealing that discovery of full katanin function can require double KO models.

We identify KATNA1/KATNAL1 as essential regulators of metaphase and anaphase spindle structure, metaphase chromosome alignment, anaphase chromosome segregation and, KATNAL1 as a cytokinesis regulator during male meiosis. While a role for katanin A-subunits in male mammalian meiosis has never previously been shown, we have previously shown the B-subunit, KATNB1 is essential for normal spindle architecture and function, and for cytokinesis in mouse spermatocytes^11,12^. In strong support of these overlapping functions being due to KATNB1 working in partnership with KATNA1 and KATNAL1, KATNB1-KATNA1 and KATNB1-KATNAL1 complexes are present throughout mouse male meiosis wherein they localize to spindle MTs^11^. That single GCKO of KATNAL1, but not KATNA1, is sufficient to cause meiotic defects, agrees with our previous data showing KATNB1-KATNAL1 complexes are the most abundant type of katanin complex during male meiosis^11^.

The misalignment of metaphase chromosomes, followed by uneven anaphase segregation in *Katnal1^GCKO/GCKO^* and *Katna1/al1^GCKO/GCKO^* meiosis is consistent with KATNA1/KATNAL1 regulating MT-chromosome attachment and/or regulating spindle dynamics via metaphase poleward ‘flux’ (reviewed by Rogers et al.^27^) and anaphase ‘Pacman-flux’ (reviewed by Rath and Sharp^28^). While future studies will need to directly test this in meiosis, katanin-mediated MT severing has been shown to directly contribute to metaphase spindle flux in HEK293 mitotic cells^29^ and Pacman-flux in *Drosophila* mitotic cells^30^.

The role for KATNAL1 in cytokinesis identified herein aligns with evidence that midbody disassembly in mitosis requires MT severing (reviewed by McNally et al.^2^ and for additional data see Elad et al.^31^) and our own data, regarding KATNB1 function in mouse male meiosis^11,12^. That only a subset of germ cells fails cytokinesis in the absence of KATNAL1 or KATNB1, suggests that other MT severing enzymes must also be involved in this process. Indeed, we have recently shown the related MT severing enzyme, spastin, also contributes to cytokinesis during male meiosis^32^.

Our data establishes KATNA1 and KATNAL1 as novel regulators of Golgi organisation and pro-acrosomal vesicle trafficking and fusion during acrosome biogenesis. This contributes to an emerging body of evidence that MT-severing enzymes regulate vesicle-based transport in spermiogenesis, as we have recently shown spastin and KATNB1 are also essential for pro-acrosomal vesicle transport and fusion events^11,32^. The contribution of KATNA1/KATNAL1 to these processes may be as simples as a role in pruning the MT tracks along which Golgi membranes are spatially positioned and pro-acrosomal vesicles are delivered. However, the interaction of KATNA1, KATNAL1 and KATNB1 with multiple proteins involved in Golgi organisation and vesicle trafficking suggests a more direct role and raises the possibility MT-severing can modulate cargo-MT interactions.

We have previously shown KATNAL2 is essential for sperm tail development via basal body-plasma membrane docking and the initiation of spermatid axoneme extension^5^. Here, we show KATNAL1 is also required, but in the subsequent assembly, and potential stability, of axoneme MT doublets, and in mitochondrial sheath and ODF development. Moreover, while KATNAL1 is not required for basal body-membrane docking, it is required for HTCA formation/integrity. The precise mechanism of KATNAL1 contribution to these processes remains an outstanding question. One possibility is that KATNAL1-mediated MT severing directly amplifies and regulates MTs for axoneme doublet formation and HTCA fortification. However, the variability in phenotypes (e.g. missing MT doublets in some tails and supernumerary MT doublets in others), is more suggestive of mis-regulated cargo trafficking. Of note, and consistent with the co-incident manchette defects in KATNAL1 loss, sperm tail components including the ODFs, and proteins required to strengthen the HTCA appear to be delivered via the MTs of the manchette, and abnormal sperm tail and manchette phenotypes often occur in parallel (reviewed by Lehti and Sironen^33^). It is thus possible that the abnormal manchette phenotypes in *Katnal1^GCKO/GCKO^* mice underpin at least some of the sperm tail defects.

The manchette phenotypes identified in the *Katnal1^GCKO/GCKO^* mice establish roles for KATNAL1 in manchette migration down the sperm nucleus, which requires ‘unzipping’ of rod-like manchette-nuclear linkers, and in controlling manchette MT length and in manchette MT disassembly. These functions are consistent with a direct MT-severing mechanism. Strikingly, the *Katnal1^GCKO/GCKO^* mouse manchette phenotypes, mirror not only those in KATNB1 loss of function mice^11,12^, but also those in KATNAL2 loss^5^. These identical yet non-redundant KATNAL1 and KATNAL2 phenotypes suggest the two proteins are executing similar functions in the manchette but severing different MT subpopulations. This is consistent with our proteomics data, demonstrating KATNAL2 is not a binding partner of KATNAL1.

Similar to the established roles of Sertoli cell KATNAL1 is required to maintain round spermatid-Sertoli cell adhesion^13^. We show germ cell KATNAL1 is involved in spermatid retention and elongated spermatid release via spermiation. Data is consistent with KATNAL1 severing the cytoskeletal connections tethering elongating spermatids to the Sertoli cell. The precise cytoskeletal targets of this action need to be identified, but of note, our IP revealed katanins interact with multiple ARP2/3 complex proteins. The ARP2/3 complex appears to facilitate spermiation, *albeit* via the disassembly of the Sertoli cell actin cytoskeleton ^19^. Of note spermiation defects in KATNAL2 mutant mice ^5^, and spermatid adhesion and spermiation defects seen in the KATNB1 mutant mice^11,12^ phenocopy those seen in *Katnal1^GCKO/GCKO^* mice. This suggests that KATNAL1-KATNB1 complexes along with KATNAL2-KATNB1 complexes facilitate spermatid release.

Finally, we defined the testis interactome for KATNA1, KATNAL1 and KATNB1. While a katanin interactome has previously been defined in HELA cells^4^, this is the first *in vivo* tissue wide interactome for each protein. A number of protein classes were identified that we predict are related to the activation/inhibition of katanin function (e.g. metabolite interconversion enzymes). Of proteins that we predict are involved specific testis functions of the katanin enzymes, cytoskeletal and vesicle-transport related proteins emerged as key protein classes. Future studies will need to define precise nature of these interactions, however they paint a picture of katanin action involving a complex crosstalk between different cytoskeletal networks notably between actin and MT networks, and suggest a direct role for katanins in regulating the interaction of dynein- and vesicle-based cargo machinery with MTs.

Collectively, this study establishes KATNA1 and KATNAL1 as novel players in male germ cell MT regulation. The data defined herein, taken together with our previous data on the roles of KATNB1^11,12^, KATNAL2^5^ and Sertoli cell KATNAL1^13^, allows us to define a model of the complete katanin ‘toolbox’ during mammalian spermatogenesis (Fig 7). This data unequivocally shows the emergence of multiple katanin A-subunits in higher eukaryotes, not only allows for functional sub-specialization and thus customized cutting of discrete MT subpopulations, but also for the protection of katanin function in critical processes such as meiosis through gene redundancy.

**Figure 7:**
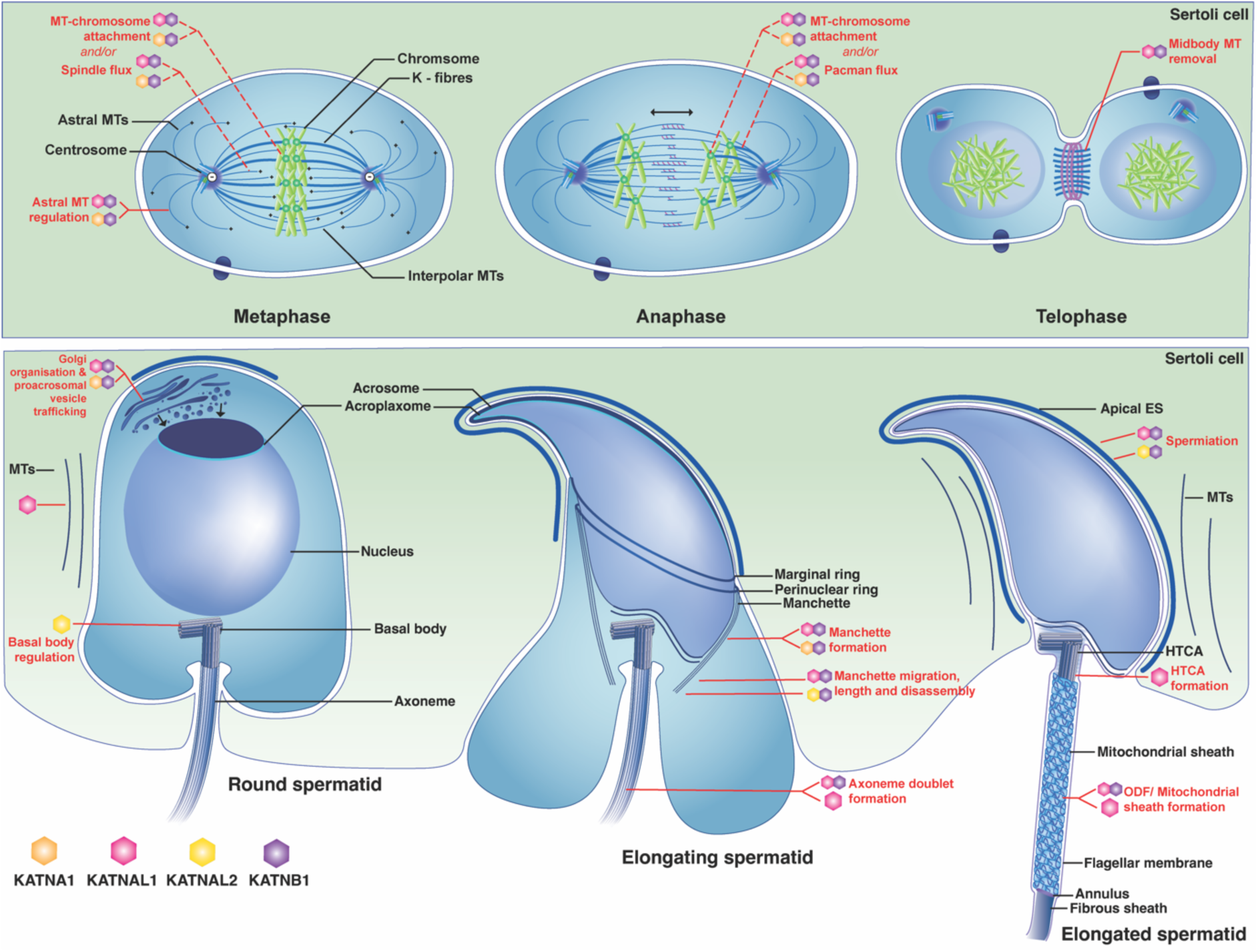
Proposed model of katanin regulation of spermatogenesis. KATNB1-independent and -dependent functions of each katanin A-subunit during male germ cell development are depicted based on the data defined herein and defined for KATNAL1 Sertoli cell function in ^13^, for KATNAL2 in ^5^ and KATNB1 defined in ^11,12^. A shared line indicates two complexes have redundant/compensatory function in the process indicated. HTCA = head to tail coupling apparatus, MTs = microtubule, ODF = outer dense fibers.

## Methods

### Mouse model production

All animal procedures were approved by the Monash University School of Biological Sciences Animal Experimentation Ethics Committee and the University of Melbourne Ethics committee and conducted in accordance with National Health and Medical Research Council (NHMRC) Guidelines on Ethics in Animal Experimentation.

To the generate *Katna1^Flox/Flox^* mice, *Katna1* KO-first conditional-ready mice (*Katna1^KO/WT^*) were generated through the Monash University Node of the Australian Phenomics Network using a EUCOMM *Katna1* KO-first conditional-ready ES clone (EPD0586_1_B10) as described^34^. To disrupt the *Katna1* gene, the FRT-LacZ-loxP-Neo-FRT-loxP-*Katna1*-*exons6*-*7*-loxP cassette was inserted into intron 5 of the *Katna1* gene (Fig S1C). To generate conditional *Katna1* mouse (*Katna1^Flox/Flox^*), mice heterozygous for the knockout-first allele (*Katna1^KO/WT^*) were then crossed with a transgenic line carrying the *Flp* recombinase gene. This resulted in removal of the *FRT* flanked gene-trap cassette, exposing a floxed *Katnal1* exon 6-7 and reverting mice to a wild-type phenotype. *Katna1^GCKO/GCKO^* were then generated using a two-step breeding strategy. (1) *Katna1^Flox/Flox^* females were mated with a *Stra-8-cre* transgenic line to produce *Katna1^WT/Flox,Stra8-cre+^* males. (2) *Katna1^WT/Flox,Stra8-cre+^* males were mated with *Katna1^Flox/Flox^* females to produce *Katna1^Flox/Flox,Stra8-cre+^* progeny (*Katna1^GCKO/GCKO^*). In these mice, gene disruption relied on Cre-lox recombination mediated excision of *Katna1* exons 6 and 7 (ENSMUSE00001254109, ENSMUSE00001247569), and the *Stra8-Cre* used is expressed from late spermatogonia onwards ^35^.

*Katnal1^Flox/Flox^* mice, wherein exon 3 (ENSMUSE00001360953) of the *Katnal1* gene is flanked by loxP sites, were generated by the Monash Genome Modification Platform using CRISPR/Cas9 technology. Guide RNA target sites flanking the target exon (5’ end of exon - 5’ TCAATGTCTGAGGTGGATAA 3’ and 3’ end of exon - 5’ CAAAGAAGTCTTTTAGACCC 3’) were identified using the UCSC Genome Browser. A single stranded DNA homology directed repair template, was generated from a *Katnal1* conditional KO targeting construct using the Guide-it Long ssDNA Strandase Kit (Takara, 632645). The following oligonucleotides were used to generate a PCR product from which the ssDNA was generated: 5’/5Biosg/GATGCTTAGCACTGGCTGG 3’ and 5’ GAACGTGTGCTATTTGGTCTTTAAG 3’. The CRISPR/Cas9 ribonucleoprotein complex and single stranded DNA homology repair templates were microinjected into C57BL/6J zygotes at the pronuclei stage and transferred into the uterus of pseudo-pregnant females. *Katnal1^GCKO/GCKO^* mice were generated by intercrossing *Katnal1^Flox/Flox^* mice with a *Stra8-cre* transgenic line as above. *Katnal2^Flox/Flox^* mice were generated as previously described^5^. *Katna1/al1^Flox/Flox^* and *Katna1/al2^Flox/Flox^* mice were generated by breeding *Katna1^Flox/Flox^* mice with either *Katnal1^Flox/Flox^* or *Katnal2^Flox/Flox^* mice, respectively. Double germ cell deletion was then achieved using by intercrossing double floxed mice with a *Stra8-cre* transgenic line as above.

All mice were maintained on a C57BL/6 background. To control for background strain effects on phenotype, *Katna1^Flox/Flox^*, *Katnal1^Flox/Flox^*, *Katna1/al1^Flox/Flox^* and *Katna1/al2^Flox/Flox^* mice mice were used as controls for *Katna1^GCKO/GCKO^*, *Katnal1^GCKO/GCKO^*, *Katna1/al1^GCKO/GCKO^* and *Katna1/al2^GCKO/GCKO^* mice, respectively. Mouse genotypes were determined from tail biopsies using real time PCR with specific probes designed for each allele (Transnetyx, Cordova, TN). Efficiency of exon excision in *Katna1^GCKO/GCKO^*, *Katnal1^GCKO/GCKO^*, *Katna1/al1^GCKO/GCKO^* and *Katna1/al2^GCKO/GCKO^* mice, was determined by qPCR of isolated germ cells as described below. Protein levels were determined by western blotting of germ cell and testis lysates as described below.

### Quantitative qPCR

Purified germ cells were prepared via the STAPUT method as detailed in ^36^. Total RNA was then extracted from purified germ cells using Trizol (Life Technologies), and reverse-transcribed into cDNA using SuperScript III Reverse Transcriptase (Life Technologies). Primers spanning *Katna1* exons 6 and 7 (5’-TCAGGAAGCAGTGGTGTTACC-3’ forward and 5’**-**AAGGGTCTTTCCAGTGCCAG-3’ reverse), *Katnal1* exons 3 and 4 5’- CCGTGTCTTGCCGAGATGAA-3’forward and 5’-GCGATTTGGACGCCTGATTT-3’ reverse) and *Katnal2* exons 2 and 3 (5’-AAGGCTTTGGAGGAGGAGAC-3’ forward and 5’**-** GGGGCCTTCTTAACCACTTT-3’) were used to verify gene deletion in the corresponding mouse models. All qPCRs were performed using SYBR Select Master Mix (Applied Biosystems). Each reaction was performed in triplicate using the Agilent Mx3000P qPCR system or an Applied Biosystems QuantStudio 3 Real-Time PCR System with the parameters of 50°C for 2 minutes, 95°C for 10 minutes followed by 40 cycles of 95°C for 15 seconds and 60°C for 1 minute. *Ppia* was amplified at the same time as an internal control (using 5’- CGTCTCCTTCGAGCTGTTT-’3 forward and 5’-CCCTGGCACATGAATCCT-’3 reverse primers), and all results were normalized to *Ppia* expression. Differential expression was analysed using the 2ΔΔ^CT^ method^37^.

### Male fertility characterisation

Male fertility of all mouse models was characterised as per^38^. Male fertility was assessed by mating 10-12 week old males with wild-type females (≥6 weeks of age). Copulatory plugs were monitored to confirm successful mating and the number of pups born per copulatory plug was recorded. Each male was test mated with 2-3 females and pup numbers averaged. Testis daily sperm production (DSP) and total epidydimal sperm content were determined using the Triton X-100 solubilisation method^39^ (n*≥*3/genotype) as modified in^11^. To assess sperm motility, sperm were backflushed as per^40^ from cauda epididymides and motility was assessed using a computer assisted sperm analyser, as described previously^41^.

For histology, testis and epididymal tissue were fixed in Bouin’s fixative and histology assessed using standard methods. Male reproductive tract histology was visualised using periodic acid-Schiff’s (PAS) and haematoxylin staining, and air-dried caudal epididymal sperm smears were visualised using haematoxylin and eosin staining (n*≥*3/genotype). Germ cell apoptosis was evaluated by immunostaining for cleaved Caspase 3 and 7 as per^42^ and as described below. The number of Caspase-positive cells in a minimum of 50 randomly selected seminiferous tubules per mouse were counted and statistical analysis conducted as described below and as per^11^(n*≥*3 mice /genotype).

### TEM

For analysis of seminiferous tubule ultrastructure, partially decapsulated testes and cauda epididymides were fixed with 5% glutaraldehyde/4% paraformaldehyde/0.02% picric acid in 0.1 M sodium cacodylate buffer, pH 7.4. as per^11^. Samples were *en bloc* stained with 2% uranyl acetate and all samples were embedded into Epon resin using standard procedures. Ultrathin sections were cut on a Leica EM UC7 Ultramicrotome and placed on Copper 100×100 square grids (ProSciTech). Sections were analysed using a FEI Talos L120CI TEM in the Ian Holmes Imaging Centre (The University of Melbourne, Parkville)

### Antibody use

Primary antibodies used included those against acetylated-tubulin (T6793, Sigma, RRID: Ab_477585, ascites fluid, 1 in 10,000), *α*-tubulin (T5168, Sigma, ascites fluid, 1 in 50,000), cleaved-Caspase 3 (9664, Cell Signalling, 0.5 μg ml^-1^), cleaved-Caspase 7 (9491, Cell Signalling, 0.23 μg ml^-1^), KATNA1 (as described previously ^11^), KATNAL1 (as described previously^11^), KATNB1 (as described in ^11^), SUN5 (ProteinTech 17495-1-AP 0.8 µg/ml) and TUBB3 (Sigma-Aldrich, SAB3300047, 0.15 μg ml^−1^). Secondary antibodies included Alexa Fluor 488 donkey anti-mouse (A212062 Invitrogen), Alexa Fluor 488 donkey anti-rabbit (A21206, Invitrogen), and Alex Fluor 555 donkey anti-rabbit (A31572, Invitrogen). EnVision+ Dual Link System-HRP (K4063, Dako) was used to amplify signal. The specificity of immunolabelling was determined by staining sections in parallel in the absence of primary antibody or where available KO mouse tissue.

### Western blotting

Proteins were extracted from STAPUT purified germ cells ^36^, and immunoblotting conducted as previously described^36^. Images were captured using a ChemiDoc MP Imaging System (BioRad). The molecular weights of detected proteins were estimated using a PageRuler Plus Prestained Protein Ladder (ThermoFisher) and the Image Lab 6.0 Molecular Weight Analysis Tool (BioRad).

### Immunochemistry and imaging

To assess KATNA1 testis localisation, 5 μm cryostat testis sections were fixed in 1:1 methanol-acetone and permeabilised in 0.2% Triton X-100 in PBS as per^43^. To assess testis localisation of all other proteins, Bouin’s fixed testes were processed into paraffin and 5 μm sections prepared using standard methods. Paraffin sections were dewaxed and heat-mediated antigen retrieval was conducted by microwaving sections 10 mM citrate buffer (pH 6) for 16 minutes as per ^44^. For sections to be used for chromogenic immunohistochemistry (*α*-tubulin, acetylated-tubulin, cleaved-caspase 3 and 7), endogenous peroxidase activity was then blocked via incubation with 5% H_2_O_2_ for 5 minutes. For all testis immunolabelling, non-specific protein binding was minimised by incubation with CAS-block (Invitrogen) for 30 minutes and sections were then incubated with primary antibodies diluted in DAKO Antibody Diluent overnight at 4°C. For immunofluorescent labelling, sections were incubated with the appropriate Alexa Fluor secondary antibodies diluted 1in 500 for 1 hour at room temperature. DNA was then visualised using 1 µg ml^-1^ 4’,6-diamidino-2-phenylindole (DAPI, Invitrogen) or 4 µM TO-PRO-3 Iodide (TOPRO, Thermo Scientific). Acrosomes were visualized using 0.5 µg ml^-1^ lectin peanut agglutinin (PNA), Alexa Fluor 488 conjugate (L21409, Life Technologies). For chromogenic immunohistochemistry, sections were incubated with DAKO EnVision+ Dual Link System-HRP for 30 minutes at room temperature. The Liquid DAB+ Substrate Chromogen System (DAKO, K3468) was then applied as per the manufacturer’s instructions. Nuclei were counterstained using Mayer’s haematoxylin (Amber Scientific), followed by Scott’s tap water (Amber Scientific).

To assess SUN5 localisation in cauda epididymal sperm, sperm were washed in PBS and airdried onto slides. Sperm were then fixed in 4% paraformaldehyde for 10 minutes. Sperm were then permeabilised, blocked for non-specific protein binding and immunofluorescently labelled as above.

Immunofluorescent images were collected using a Leica TCS SP8 confocal microscope (Leica Microsystems) at the Monash Micro Imaging (Monash University) and the University of Melbourne Biological Optical Microscopy Unit (University of Melbourne, Parkville). A 63x/1.40 HC PL APO CS2 oil immersion objective was used to take all immunofluorescent images. Bright field images were collected cellSens Imaging Software (Olympus) on an Olympus BX53 light microscope equipped with an Olympus DP80 camera.

### IP-MS

For IP assays, wild-type mouse testes (n=3 per IP assay) were lysed in 1% NP-40 (PBS) supplemented with protease inhibitor cocktail (Calbiochem, 539134). KATNA1, KATNAL1 and KATNB1 and their binding partners were then co-immunoprecipitated using the Pierce Co-immunoprecipitation Kit (Thermo Scientific, 26149) as per the manufacturer’s instructions, in combination with either our KATNA1 (10μg per IP column), KATNAL1 (5μg) or KATNB1 (5μg) inhouse antibody. Briefly, to reduce non-specific binding, testis protein lysates were first pre-cleared by incubation with 400μl of Pierce control agarose resin slurry for a 0.5-1 hour at 4°C with gently rocking. For each pre-cleared testis lysate, 5 mg was then loaded into two IP columns, one which contained the target antibody covalently coupled to the amine reactive Pierce AminoLink Plus Coupling Resin and one which contained the target antibody and a non-amine reactive Pierce Control Agarose Resin (negative control), and incubated overnight at 4°C. Lysate flow through was removed, columns were washed and bound protein eluted and collected as per the manufacturer’s instructions. The eluted proteins were neutralised with 1.5M Tris-HCL (pH 8.8), reduced with TCEP, then digested by incubation with trypsin overnight at 37°C. Peptide samples were freeze dried and resuspended in 2% acetonitrile (ACN)/0.1% formic acid (FA) (diluted in MilliQ water) before concentrated FA was added to adjust the pH to <3. Samples were purified using reverse-phase chromatography C18 stage tips, which aim to desalt and fractionate in-solution peptides at an acidic pH. The tips were initially activated with 100% methanol and equilibrated with 0.1% FA, before samples were loaded into them and desalted with 0.1% FA. Peptides were eluted with 80% ACN/0.1% FA, freeze dried, resuspended in 2% ACN/0.1% FA and sonicated for 15 minutes in a water bath.

The purified peptide samples were analysed via nano liquid chromatography coupled to tandem mass spectrometry (LC-MS/MS). Briefly, the KATNA1 IP and KATNB1 IP samples were analysed at the Monash Proteomics and Metabolomics Facility (Monash University) using an Orbitrap Fusion Tribrid mass spectrometer (ThermoFisher, USA) and a Q-Exactive Hybrid Quadrupole - Orbitrap mass spectrometer (ThermoFisher, USA), respectively. Both instruments were coupled to an UltiMate 3000 RSLCnano system (ThermoFisher, USA) equipped with an Acclaim PepMap RSLC C18 nanoViper Analytical Column (100Å, 75 μm X 50 cm, 2 μm; ThermoFisher, USA) and an Acclaim PepMap 100 C18 HPLC nanoViper Trap Column (100Å, 100 μm X 2 cm, 5 μm; ThermoFisher, USA). The tryptic peptides were separated by increasing concentrations of 80% acetonitrile (ACN) / 0.1% formic acid at a flow rate of 250 nl/min for 128 min and 158 min, respectively. Both mass spectrometers were operated in data-dependent acquisition (DDA) mode to automatically switch between full scan MS and MS/MS acquisition. Each survey full scan (m/z 375–1800) in the Orbitrap Fusion Tribrid acquisitions was acquired in the Orbitrap with 120,000 resolution (at m/z 200) after accumulating ions to a Normalised AGC Target of 250% with a maximum injection time of 54 ms. Dynamic exclusion was set to 15 seconds. Adhering to a 2 sec cycle time, the most intense multiply charged ions (z ≥ 2) were sequentially isolated and fragmented in the collision cell by higher-energy collisional dissociation (HCD) and ms2 scans were acquired with a fixed injection time of 54 ms, 30,000 resolution and a Normalised AGC Target of 400%. Very similar parameters were used for the Q-Exactive Hybrid Quadrupole – Orbitrap acquisitions.

The KATNAL1 IP purified peptide samples were analysed at the University of Melbourne Mass Spectrometry and Proteomics Facility, using an Orbitrap Eclipse Tribrid Mass Spectrometer (ThermoFisher, USA) equipped with a nano ESI interface coupled to an Ultimate 3000 nano HPLC (Thermo Fisher, USA). The LC system was equipped with an Acclaim Pepmap 100 C18 HPLC nanoViper Trap Column (100 Å, 75 μm X 2 cm, 3 μm; ThermoFisher, USA) and an analytical column as above. The enrichment column was injected with the tryptic peptides (1 µL, 100 ng) at an isocratic flow of 5 μL/min of 2% *v/v* CH_3_CN containing 0.05% *v/v* trifluoroacetic acid for 6 min, applied before the enrichment column was switched in-line with the analytical column. The eluents were 0.1% v/v formic acid (solvent A) in H_2_O and 100% *v/v* CH_3_CN in 0.1% *v/v* formic acid (solvent B). The gradient was at 300 nl min^-1^ from (i) 0–6 min, 3 % B; (ii) 6–35 min, 3–23 % B; (iii) 35-45 min, 23-40 % B; (iv) 45-50 min, 40-80 % B; (v) 50-55 min, 80–80 % B; (vi) 55-55.1 min, 80-3 % B; (vii) 55.1-65 min, 3-3 % B. The Eclipse Orbitrap mass spectrometer was operated in the data-dependent mode, wherein full MS^1^ spectra were acquired in a positive mode over the range of *m/z* 375-1500, with spray voltage at 1.9kV, source temperature at 275 °C, MS^1^ at 120,000 resolution, normalized AGC target of 100 % and maximum IT time of 22 ms. The top 3 second method was used and selecting peptide ions with charge states of ≥ 2-7 and intensity thresholds of ≥ 5E4 were isolated for MS/MS. The isolation window was set at 1.6 *m/z*, and precursors were fragmented using higher energy C-trap dissociation (HCD) at a normalised collision energy of 30, a resolution of 15,000, a normalized AGC target of 100% and automated IT time.

The raw MS data were analysed with the MaxQuant software suite for the identification and quantification of peptides/proteins from the Mouse SwissProt database. Fixed modifications of carbamidomethylation of cysteine as well as variable oxidation of methionine and protein N-terminal acetylation were considered. Trypsin/P was set as the protease with a maximum of 2 missed cleavages. Statistical analysis was conducted with the Perseus software considering protein and PSM false discovery rates (FDR) both set at < 0.01. For each IP assay, the majority UniProt ID(s) for each significantly enriched protein group were then analysed using PANTHER Gene Ontology and protein classification^15^. All MS proteomics data have been deposited to the ProteomeXchange Consortium via the PRIDE partner repository with the dataset identifier PXD038404 ^45^.

### Statistics

Statistical analysis of germ cell apoptosis data was conducted in R version 3.5.1^46^. Generalised linear mixed (GLM) models were used to compare the number of caspase-positive cells per tubule across genotypes, with animal ID included as a random effect to account for repeated measures per individual. For each model, Akaike information criteria (AIC) estimates were used to select the most appropriate error distribution and link function (i.e. poisson, negative binomial, zero-inflated poisson, zero-inflated negative binomial) using the glmer function (lme4 package^47^) and glmmTMB function (glmmTMB, package^48^). For all models, a zero-inflated negative binomial distribution (fitted with glmmTMB, using the ziformula argument) was selected as the most appropriate error distribution and link function (i.e. had the lowest AIC score).

All other statistical analysis, other than for the proteomics (described above), was conducted in Graphpad Prism 9.0. The statistical significance of differences between two groups was determined with an unpaired Student t-test. Significance was defined as a P value of <0.05.

## Acknowledgements

We thank Dr Liza O’Donnell and Dr Duangporn Jamsai for advice on the initial stages of the project. We thank Ms G. Gemma Stathatos, Ms Lauren Rogers and Ms Georgia Shering for technical assistance. We thank Associate Professor Nicholas Williamson, Dr Ching-Seng Ang and Dr Michael Leeming for technical assistance and advice regarding the MS analysis. We thank the following platforms for the use of their facilities and services: at The University of Melbourne (Parkville), the Biological Optical Imaging Platform, the Ian Holmes Imaging Centre, the Mass Spectrometry and Proteomics Facility, the Melbourne Bioresources Platform and the Melbourne Histology Platform; and at Monash University (Clayton), the Monash Animal Research Platform, the Monash Gene Modification Platform, the Monash Histology Platform, Monash Micro Imaging and the Monash Proteomic and Metabolomic Facility.

## Author Contributions

Conceptualisation, J.E.M.D, M.K.O; Methodology, J.E.M.D, A.E.O, D.J.M, R.B.S, B.J.H and M.K.O; Formal Analysis, J.E.M.D, M.G, K.W, A.E.O, D.J.M, J.N, R.B.S, B.J.H, M.K.O; Investigation, J.E.M.D, M.G, K.W, A.E.O, D.J.M, J.N, R.B.S, B.J.H; Writing – Original Draft, J.E.M.D, M.G, M.K.O; Writing – Review & Editing, J.E.M.D, R.B.S, M.K.O; Visualisation, J.E.M.D, M.G, K.W; Supervision, J.E.M.D, M.K.O; Project Administration, J.E.M.D, M.K.O; Funding Acquisition, M.K.O

