## Supplemental Figs 1-6 and Supplemental Table 3 for "Male mammalian meiosis and spermiogenesis is critically dependent on the shared functions of the katanins KATNA1 and KATNAL1"

**Supplemental figures**

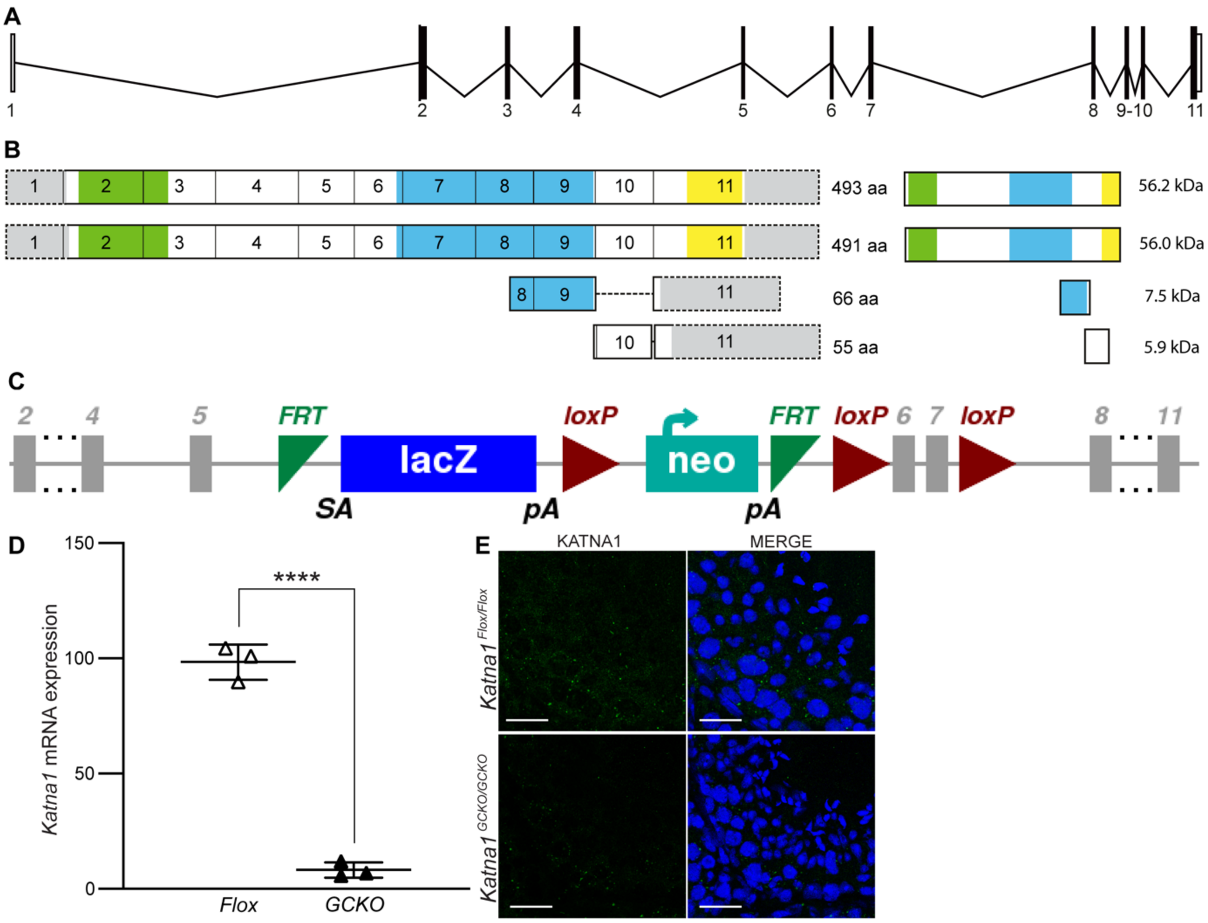


### Figure S1: Generation of the *Katna1*$^{GCKO/GCKO}$ mouse model

*Katnal1* gene (**A**) and transcript structure (**B**). The MT interacting and transport (MIT) domain, AAA ATPase domain and VPS4 domain are shown in green, blue and yellow respectively. Untranslated regions are shown in grey. (**C**) The *Katna1* KO-first conditional ready cassette. The FRT-lacZ-loxP-Neo-FRT-loxP-*Katna1* exons 6-7-loxP cassette was inserted into *Katna1* intron 5. (**D**) qPCR analysis of *Katna1* transcript levels in *Katna1*$^{GCKO/GCKO}$ (black triangles) relative to *Katna1*$^{Flox/Flox}$ (white triangles) isolated spermatocytes (n=3/genotype). ****$P$<0.0001. (**E**) *Katna1*$^{Flox/Flox}$ and *Katna1*$^{GCKO/GCKO}$ testis sections immunolabelled for KATNA1 (green). Nuclei are counterstained with DAPI (blue). Scale bars in **F** = 20 µm.

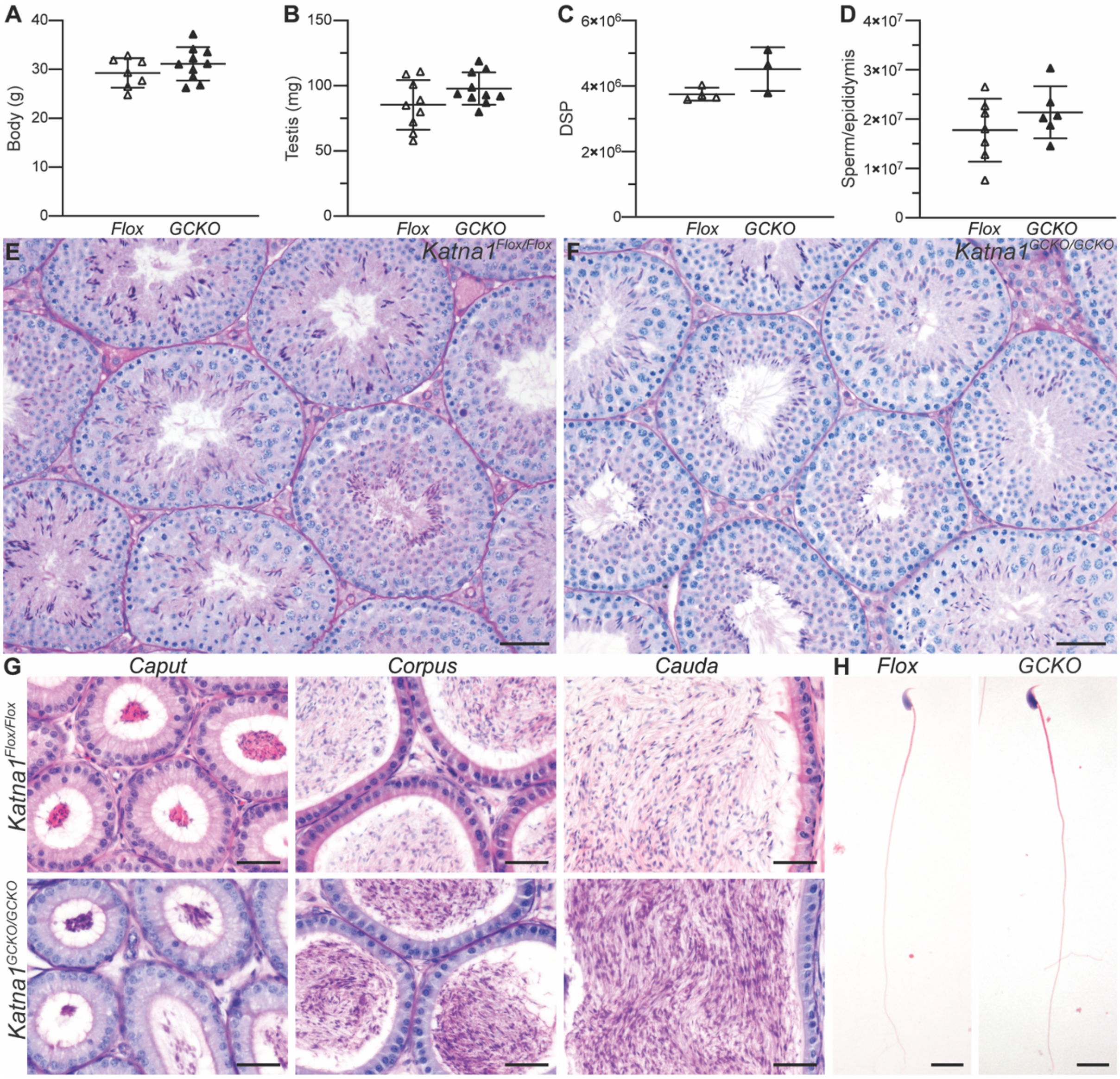


**Figure S2: KATNA1 is dispensable for male meiosis and haploid germ cell development**

Body weight (**A**), testis weight (**B**), testis DSP (**C**), and epididymal sperm content (**D**) in *Katna1*$^{GCKO/GCKO}$ mice (black triangles) and *Katna1*$^{Flox/Flox}$ controls (white triangles) (n≥3/genotype). Error bars represent mean ± SD. PAS-stained testis (**E-F**) and epididymis (**G**) sections, and hematoxylin and eosin-stained cauda epididymal sperm (**H**) from *Katna1*$^{GCKO/GCKO}$ and *Katna1*$^{Flox/Flox}$ mice. Scale bars in **E-F** = 50 μm, **G** = 40 μm and **H** = 10 μm.

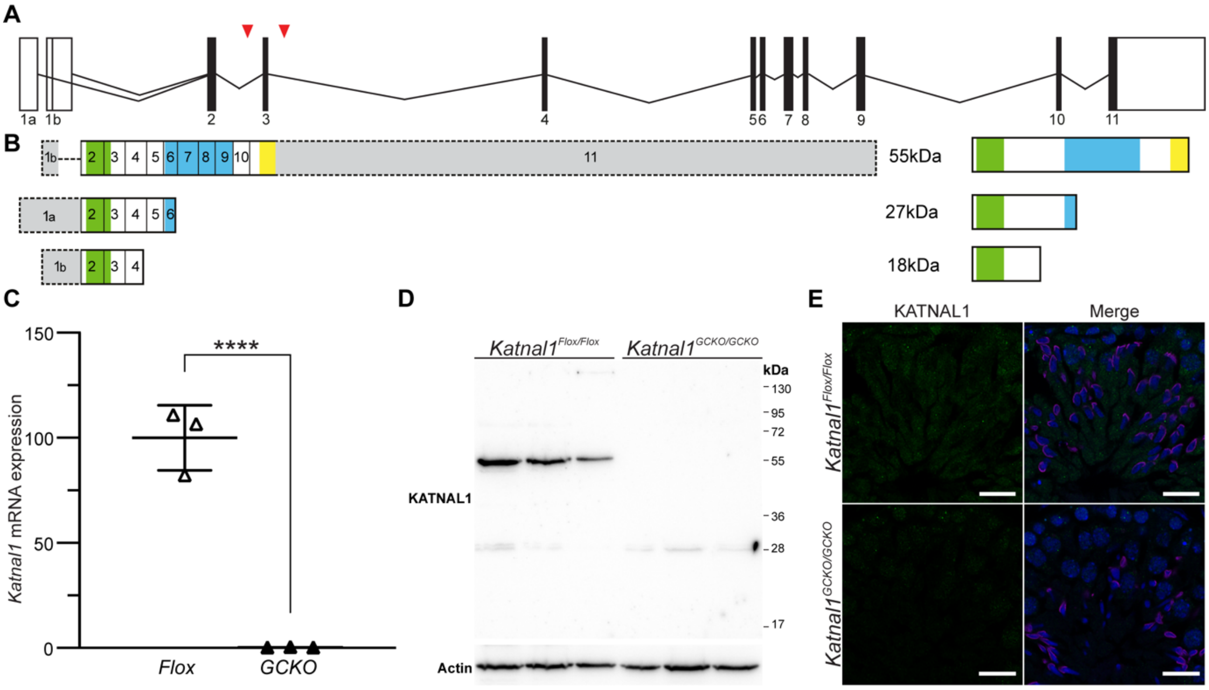


**Figure S3: Generation of the *Katnal1$^{GCKO/GCKO}$* mouse model**

*Katnal1* gene (**A**) and transcript structure (**B**). The MT interacting and transport (MIT) domain, AAA ATPase domain and VPS4 domain are shown in green, blue and yellow respectively. Untranslated regions are shown in grey. (**C**) qPCR analysis of *Katnal1* transcipt levels in *Katnal1$^{GCKO/GCKO}$* isolated spermatocytes relative to *Katnal1$^{Flox/Flox}$* controls (n=3/genotype). Data are normalised to *Ppia* and lines represent mean ± SD. ****$P$<0.0001. (**D**) Western blot analysis of KATNAL1 in isolated spermatocytes from *Katnal1$^{Flox/Flox}$* and *Katnal1$^{GCKO/GCKO}$* mice. Blots were re-probed with actin as a loading control. (**E**) Testis sections immunolabeled for KATNAL1 (green) in *Katnal1$^{Flox/Flox}$* and *Katnal1$^{GCKO/GCKO}$* mice. DNA and acrosomes were counterstained with DAPI (blue) and PNA (magenta) respectively. Scale bars = 20 µm.

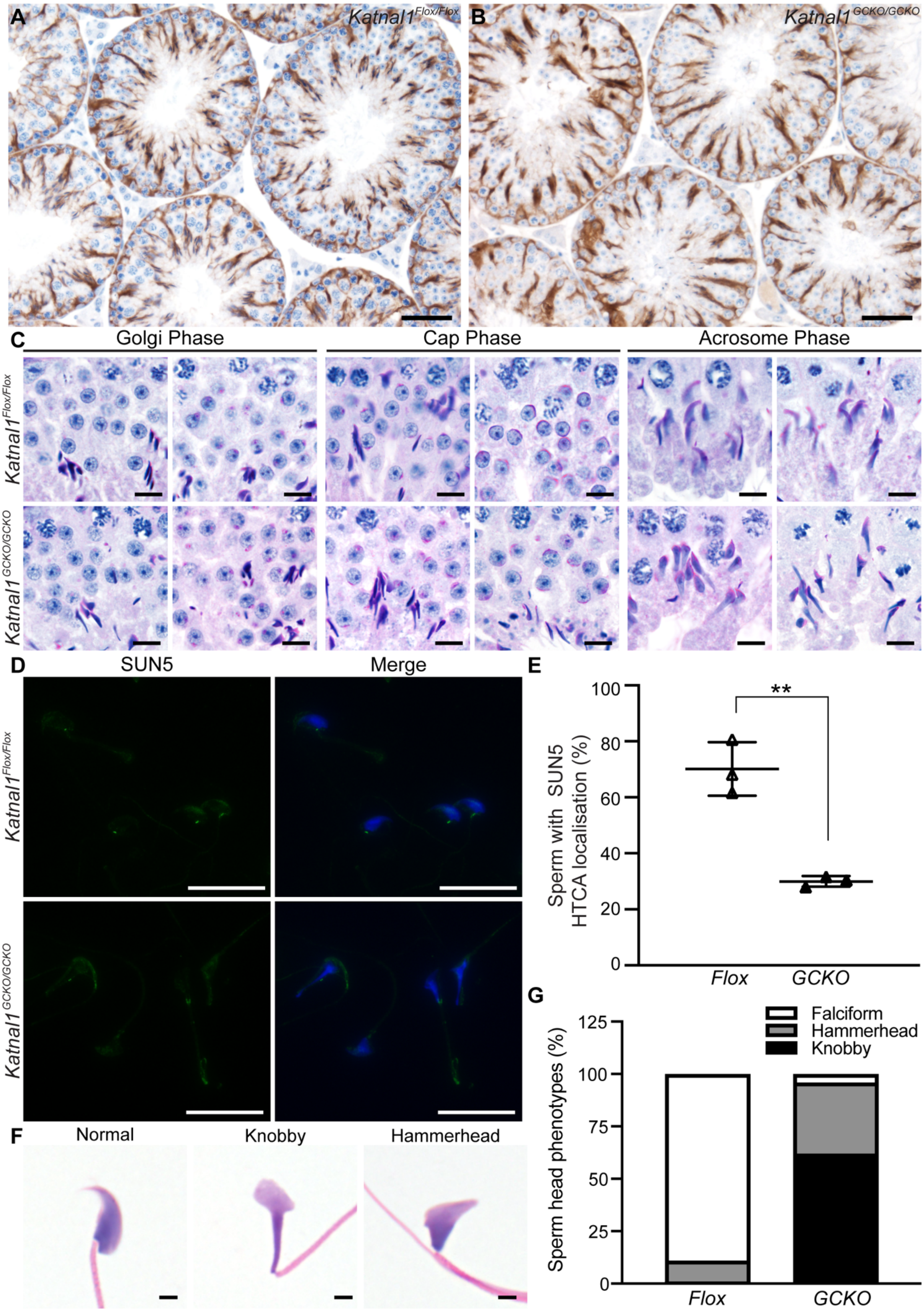


**Fig S4: *Katnal1$^{GCKO/GCKO}$* mice contain a normal Sertoli cell MT cytoskeleton and acrosome formation but have defects in sperm head shape and HTCA integrity**

*Katnal1*$^{Flox/Flox}$ (**A**) and *Katnal1*$^{GCKO/GCKO}$ (**B**) testis sections immunolabelled for the Sertoli cell specific β-tubulin isoform TUBB3. (**C**) PAS-stained testis sections showing acrosome formation in *Katnal1*$^{Flox/Flox}$ and *Katnal1*$^{GCKO/GCKO}$ mice. Progressive steps of acrosome formation are shown left to right. Roman numerals denote seminiferous tubule stage. (**D**) Representative images of epididymal sperm from *Katnal1*$^{Flox/Flox}$ and *Katnal1*$^{GCKO/GCKO}$ mice immunolabelled for SUN5 (green) as an essential component of the HTCA. Nuclei are counterstained with DAPI (blue). (**E**) Percentage of epididymal that had SUN5 localised to the base of sperm head sperm in *Katnal1*$^{Flox/Flox}$ (white triangles) and *Katnal1*$^{GCKO/GCKO}$ (black triangles) mice (n≥3/genotype, for each animal and a minimum of 100 epididymal sperm per animal). Example images of sperm head shape phenotypes (**F**) and percentage of each sperm head phenotype in in *Katnal1*$^{Flox/Flox}$ and *Katnal1*$^{GCKO/GCKO}$ mice (**G**) (n=4/genotype, for each animal a minimum of 100 epididymal sperm were assessed). $^{**}P<0.01$. Scale bars in **A** = 50 μm, **B** = 10 μm, **D** = 20 μm **F** = 2 μm.

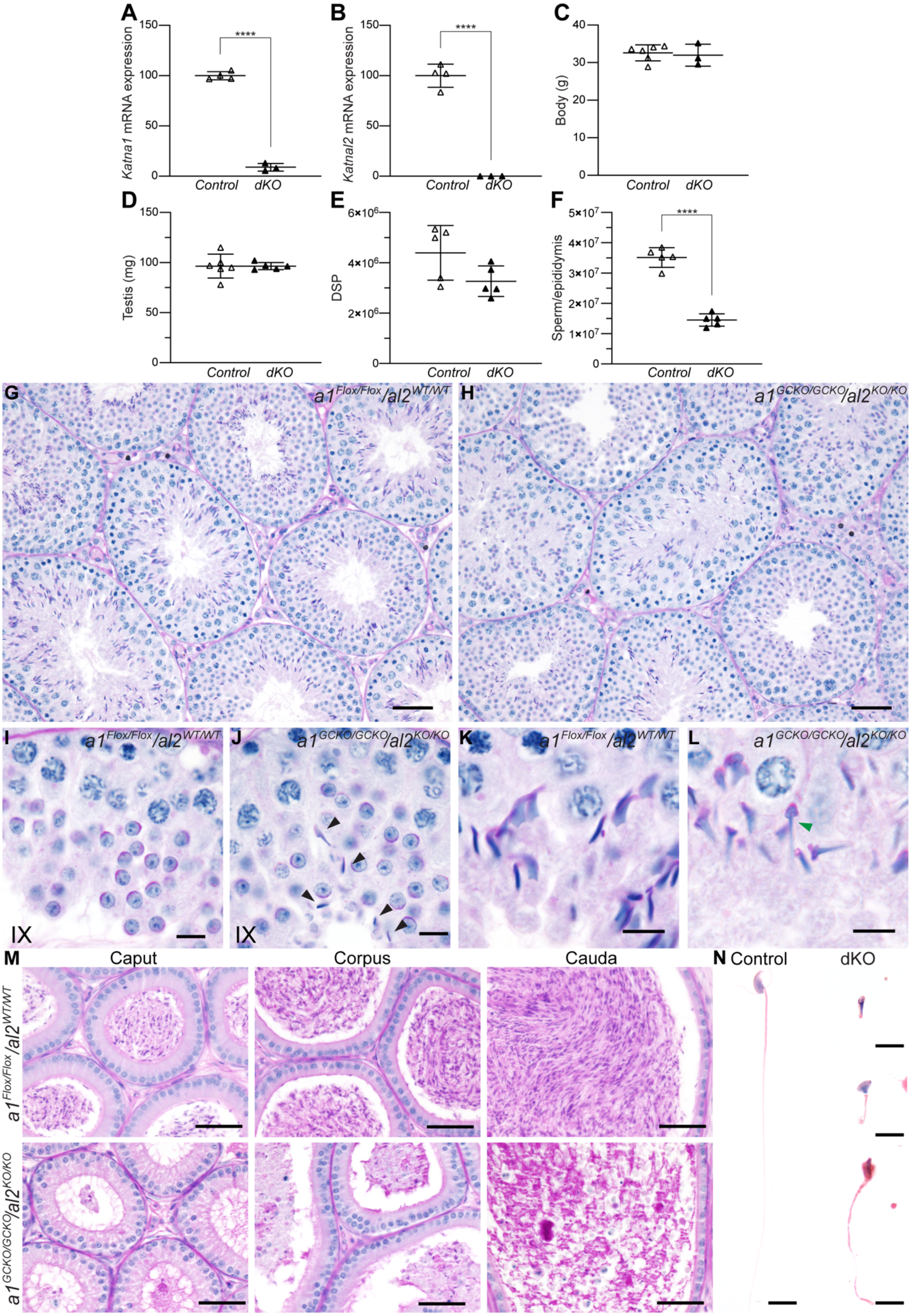
A
Katna1 mRNA expression
150
100
50
0
****
Control
dKO
B
Katnal2 mRNA expression
150
100
50
0
****
Control
dKO
C
Body (g)
40
30
20
10
0
Control
dKO
D
Testis (mg)
150
100
50
0
Control
dKO
E
DSP
6×10^6
4×10^6
2×10^6
0
Control
dKO
F
Sperm/epididymis
5×10^7
4×10^7
3×10^7
2×10^7
1×10^7
0
****
Control
dKO
G
a1 Flox/Flox/al2 WT/WT
H
a1 GCKO/GCKO/al2 KO/KO
I
a1 Flox/Flox/al2 WT/WT
IX
J
a1 GCKO/GCKO/al2 KO/KO
IX
K
a1 Flox/Flox/al2 WT/WT
L
a1 GCKO/GCKO/al2 KO/KO
M
Caput
Corpus
Cauda
a1 Flox/Flox/al2 WT/WT
a1 GCKO/GCKO/al2 KO/KO
N
Control
dKO

**Fig S5: KATNA1 and KATNAL2 do not function redundantly during mouse spermatogenesis**

qPCR analysis of *Katna1* (**A**) and *Katnal2* (**B**) transcript levels in $Katna1^{GCKO/GCKO}/al2^{KO/KO}$ (black triangles) isolated spermatocytes relative to $Katna1^{Flox/Flox}/al2^{WT/WT}$ (white triangles) isolated spermatocytes (n≥3/genotype). Body weight (**C**), testis weight (**D**), testis DSP (**E**), and epididymal sperm content (**F**) in $Katna1^{GCKO/GCKO}/al2^{KO/KO}$ mice (black triangles) compared to $Katna1^{Flox/Flox}/al2^{WT/WT}$ controls (white triangles) (n≥3/genotype). Error bars represent mean ± SD. PAS stained testis (**G-L**) and epididymis (**M**) sections, and hematoxylin and eosin-stained cauda epididymal sperm (**N**), from $Katna1^{GCKO/GCKO}/al2^{KO/KO}$ and $Katna1^{Flox/Flox}/al2^{WT/WT}$ mice. Scale bars = 50 µm in **G-H**, 10 µm in **I-L**, 40 µm in **M-N**. ****$P$<0.0001.

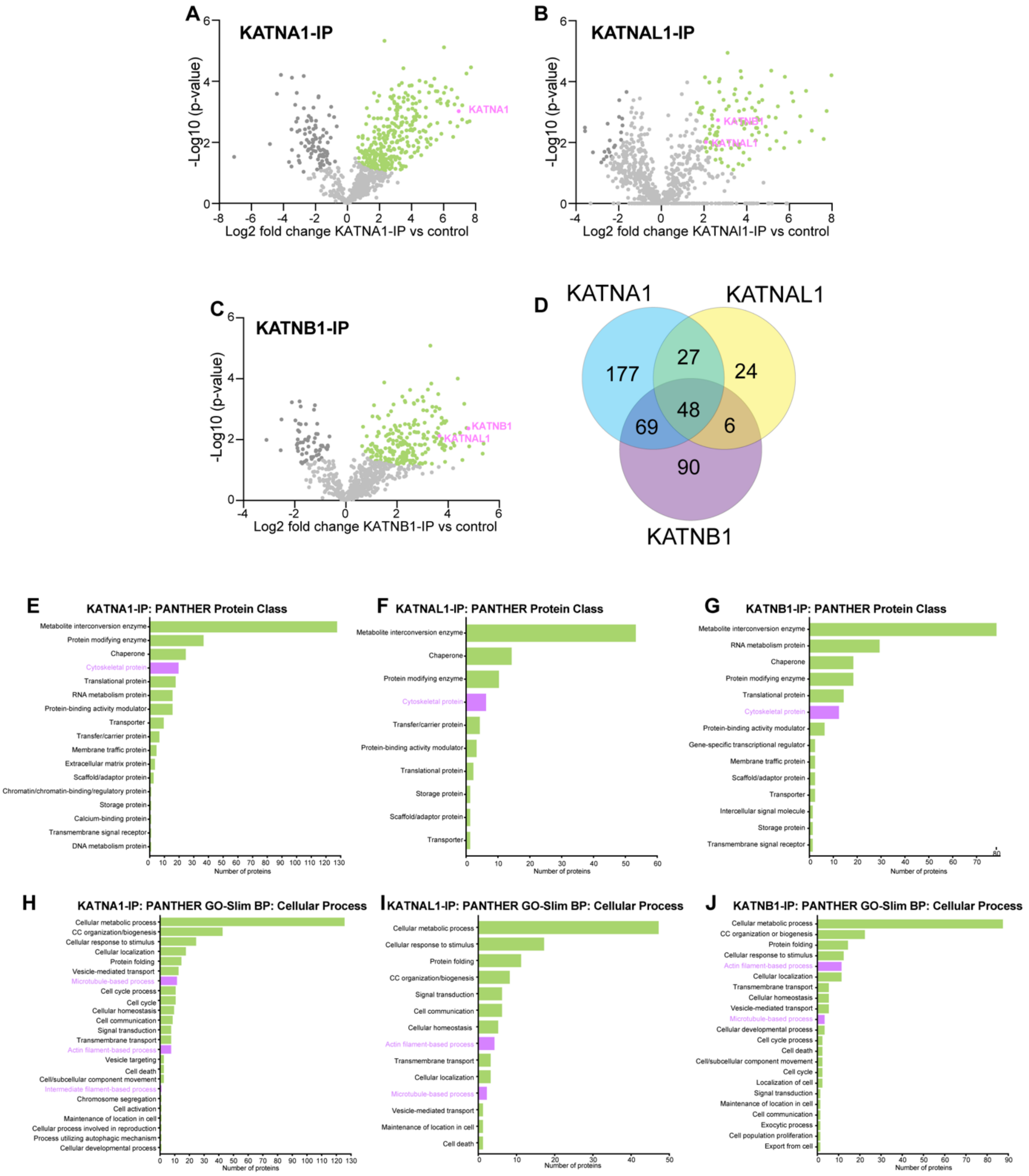


**Fig S6: Identification of KATNA1, KATNAL1 and KATNAL2 testis interaction partners**

Volcano blots showing the statistical enrichment of protein groups identified in the KATNA1 (**A**), KATNAL1 (**B**) and KATNB1 (**C**) IP-MS assays. All protein groups identified, IP-MS measurements and statistical analysis is provided in Table S1. Protein groups shown in green or pink, represent those significantly enriched in the experimental IP (KATNA1, KATNAL1 or KATNB1). For each IP-MS assay n=3 biological replicates were assessed. Venn diagram (**D**) showing the overlap in the identity of the protein groups identified in each of the KATNA1, KATNAL1 and KATNB1 IP-MS assays. PANTHER Protein Class and Gene Ontology analysis was used to analyse the proteins significantly enriched in each of the KATNA1, KATNAL1 and KATNB1 IPs (**E-J**). The PANTHER Protein Class (**E-F**) and the of the PANTHER GO-Slim Biological Process (BP) Cellular Process (GO:0009987) sub-classification (**H-J**) assigned to the proteins significantly enriched in each of the KATNA1, KATNAL1 and KATNB1 IPs. PANTHER was not able to assign all proteins identified. Detailed PANTHER Protein Class and GO-Slim analysis is provided in Table S2.

### Supplemental tables

**Table S1:** Significantly enriched proteins identified in KATNA1, KATNAL1 and KATNB1 testis co-IP experiments with subsequent MS analysis. Relates to Fig S7 and Tables S2 and S3. **Attached excel file*

**Table S2**: PANTHER analysis of proteins identified by MS as significantly enriched in either KATNA1, KATNAL1 or KATNB1 testis co-immunoprecipitates. Relates to Fig S7 and Tables S1 and S3. **Attached excel file*

**Table S3.** Functional descriptions of selected KATNA1 (A1), KATNAL1 (AL1) and KATNB1 (B1) candidate testis binding proteins of interest.

| Gene name(s) | Protein name(s) | Function | Katanin IP significantly enriched in |
|---|---|---|---|
| **Actin-related proteins** | | | |

| | | | |
|---|---|---|---|
| *Actn1/Actn4* | Alpha-actinin 1 and 4 | F-actin cross-linking proteins and part of the membrane-associated spectrin protein family. ACTN4 has been localised to the mouse spermatid manchette perinuclear ring where it binds SEPT14 [49]. | B1 |
| *Actr2 (Arp2)* | Actin-related protein 2 | Components of the ARP2/3 complex, which regulates (F-)actin filament polymerisation and mediates the formation of branched F-actin networks. Components are enriched in ectoplasmic specialisations (junctions between Sertoli cells and male germ cells), and data supports roles in blood-testis-barrier integrity, spermiogenesis and spermiation [16-19]. | B1 |
| *Actr3 (Arp3)* | Actin-related protein 3 | | AL1 |
| *Arpc2* | Actin Related Protein 2/3 Complex Subunit 2 | | A1, B1 |
| *Arpc3* | Actin Related Protein 2/3 Complex Subunit 3 | | AL1, B1 |
| *Arpc4* | Actin Related Protein 2/3 Complex Subunit 4 | | A1, B1 |
| *Arpc5* | Actin Related Protein 2/3 Complex Subunit 5 | | B1 |
| *Add1* | Alpha adducin. | Actin-binding protein which promotes assembly of the spectrin-actin network. Functions in mammalian mitotic spindle assembly (*in vitro*), by maintaining the integrity of the centrioles/spindle poles [50,51]. | A1 |
| *Capza1* | F-actin-capping protein subunit alpha-1 | F-actin-capping proteins. They bind to the barbed-ends of F-actin to block monomer exchange. | A1, AL2, B1 |
| *Capza2* | F-actin-capping protein | | A1, AL2 |

| | | | |
|---|---|---|---|
| | subunit alpha-2 | | |
| *Flna* | Filamin a | Actin binding protein that promotes orthogonal branching of F-actin and links F-actin to membrane glycoproteins. Involved in ciliogenesis [52]. | A1, B1 |
| *Gmfb* | Glia Maturation Factor Beta | Actin depolymerisation factor which binds the ARP2/3 complex to remodel branched F-actin networks. | B1 |
| *Gsn* | Gelsolin | Actin-modulating protein that caps the plus ends of actin monomers and filaments, blocking monomer exchange. Involved in F-actin nucleation and capable of severing F-actin. | A1 |
| *Myl6* | Myosin light chain 6 | Actin motor protein. Present in the mouse spermatid manchette and a testis interacting partner of SPATA6, a protein required for HTCA formation [53]. | A1 |
| *Pfn2* | Profilin 2 | Profilins bind actin monomers and promote F-actin formation. Involved in the negative regulation of endocytosis [54]. | A1 |
| *Sptan1* | Spectrin alpha, non-erythrocytic 1 | Part of the spectrin protein family. Spectrins are membrane-associated scaffolding proteins which link actin to the plasma membrane and are involved in intracellular organelle organisation. | B1 |
| *Zyx* | Zyxin | Focal adhesion component and involved in F-actin polymerisation. Present in rat Sertoli cell junctions including ectoplasmic specialisations and tubulobulbar complexes, where data suggests it is an adapter between junctions and cytoskeletal filaments (F-actin, MTs, and intermediate filament) [55]. | A1 |
| **MT-related proteins** | | | |
| *Arl3* | ADP-ribosylation factor - like protein 3 | Small GTPase that is involved in intraflagellar transport and ciliogenesis. In mice, present in the spermatid manchette and required for normal sperm head shaping and tail development [56]. | A1 |
| *Actr1a* (*Arp1*) | Alpha-centractin | Essential components of the dynactin complex. Dynactin binds dynein and MTs, to aid dynein-mediated cargo transport along MTs. Dynactin drives chromosomes segregation in C. elegans spermatocytes [57] and its disruption leads to male infertility in *Drosophila* [58]. ACTR1A localises to the Golgi apparatus, acrosome, manchette and centrioles of rat spermatids [59]. | A1, B1 |
| *Dctn2* | Dynactin subunit 2 | | A1 |
| *Dnaaf4* *(Dyx1c1)* | Dynein axonemal assembly factor 4 | Conserved role in axonemal dynein assembly and ciliary motility. Required for mammalian male fertility [60]. | A1 |

| | | | |
|---|---|---|---|
| *Mapre1 (Eb1)* | MT associated protein RP/EB family member 1 | MT plus-end tracking protein (+TIP). Regulates MT dynamics and organisation and promotes MT nucleation and elongation. Involved in mitotic spindle positioning and chromosome segregation, and negatively regulates actin nucleation in human cells. Regulates MT and actin networks in the rat Sertoli cell blood-testis-barrier (*in vitro*) [61,62]. | B1 |
| *Nudc* | Nuclear distributio n C, dynein complex regulator | Nuclear movement protein that associates with dynein-dynactin complexes, tubulin and MT organising centres (MTOCs). Established roles in human mitosis including spindle formation, chromosome segregation and cytokinesis. Role in cell proliferation in humans. | A1, AL1, B1 |
| *Tuba1b/Tuba1c/ Tuba4a/Tuba8* | Alpha-tubulin 1b, 1c, 4a and 8 | Alpha tubulin subunits of MTs. | A1 |
| *Tubb4a/Tubb4b* | Beta-tubulin 4a and 4b | Beta tubulin subunits of MTs. | A1 |
| **Intermediate filament related proteins** | | | |
| *Vim* | Vimentin | Type III intermediate filament protein, present throughout the Sertoli cell cytoplasm. | A1 |
| **Spermatogenesis related proteins** | | | |
| *Acrbp* | Acrosin-binding protein | Acrosomal protein. Required for the formation of the acrosomal granule during acrosome biogenesis and maintenance of a single acrosomal vesicle [63]. | AL1 |
| *Fabp9* | Fatty acid binding protein 9 | Member of the fatty acid binding family, which have roles in fatty acid transport. FABP9 is a component of the mouse sperm perforatoirum and perinuclear theca and is involved in sperm head shaping [64]. | A1, AL1 |
| *Hspa2* | Heat shock protein family A (Hsp70) member 2 | Molecular chaperone essential for male fertility. Loss causes meiosis arrest at the end of pachytene and mass germ cell apoptosis in mouse spermatogenesis [65]. Required for synaptonemal complex desynapsis in meiosis I prophase spermatocytes [66]. Transition protein chaperone in spermiogenesis [67]. | A1, B1 |
| *Tex15* | Testis expressed 15, meiosis and synapsis associated | Essential for male meiosis, loss in mice causes meiosis arrest before mid-pachytene. Required for synaptonemal complex formation and meiotic recombination in spermatocytes [68]. Involved in the testis piRNA pathway [69]. | A1 |

| Vesicle/membrane transport related proteins | | | |
|---|---|---|---|
| *Arf1/Arf2/Arf3* | ADP ribosylation factor 1 and 3 | Golgi apparatus associated small GTPases. Involved in Golgi vesicle budding and uncoating, intra-Golgi transport and cytokinesis. | A1 |
| *Erp29* | Endoplasmic reticulum protein 29 | Luminal ER protein, involved in ER to cis-Golgi transport. | A1, B1 |
| *Gdi2* | GDP dissociation inhibitor 2 | Negatively regulates intracellular membrane trafficking by keeping Rab proteins in their inactive state. | A1 |
| *Hspa8* | Heat shock protein family A (Hsp70) member 8 | A chaperone protein, that also mediates the ATP-dependent disassembly of clathrin-coated vesicles. | A1, AL1, B1 |
| *Lman1* | Lectin, mannose binding 1 | LMAN1 and MCFD2 complex to form a specific cargo receptor that is involved in ER to cis-Golgi protein transport. | A1 |
| *Mcfd2* | Multiple coagulation factor deficiency 2, ER cargo receptor complex subunit | | A1 |
| *Rab11fip1* | RAB11 family interacting protein 1 | Involved in the RAB11-dependent recycling of endosomal vesicles. Involved in vesicular trafficking events required for cytokinesis in mammalian cells. | B1 |
| *Rab14* | RAB14, member RAS oncogene family | Rab GTPase. Regulates Golgi to endosome transport. Present in the mouse sperm acrosome and tail. | B1 |
| *Rab2a/Rab2b* | RAB2A and 2B, member RAS | Rab GTPases. Regulate ER to Golgi transport and the compacted morphology of the Golgi. Present in the Golgi apparatus of mammalian round spermatids, and redistributed to the acrosome of elongating spermatids [70,71]. Testis interacting proteins of FAM71F1, a regulator of acrosome formation [71]. | A1 |

| | | | |
|---|---|---|---|
| | oncogene family | | |
| *Rab5a/Rab5b/ Rab5c* | RAB5A, 5B, and 5C member RAS oncogene family | Rab GTPases. RAB5 proteins are master regulators of early endosome trafficking (vesicle biogenesis, trafficking, early to late endosome maturation). Can contribute to exocytosis. Present in the acrosome of round spermatids [22]. | A1 |
| *Rab6a/Rab6b* | RAB6A and 6B, member RAS oncogene family | Rab GTPases. Regulates intra-Golgi transport, a variety of anterograde and retrograde Golgi transport pathways and the compacted morphology of the Golgi. Present in the round spermatid Golgi and is transported to the inner acrosomal/apical nuclear membrane during acrosome development[22,72]. Knockout of RAB6 interactor VSP13B, results in failure of RAB6-vesicles targeting to the nuclear membrane and a failure of acrosome development [72]. | B1 |
| *Sec31a* | SEC31 homolog A, COPII coat complex component | Component of the COPII complex which mediates formation of ER transport vesicles. | A1 |
| *Vcp (p97)* | Valosin containing protein | AAA-ATPase, which mediates intracellular membrane fusion and is involved in ER and the Golgi biogenesis. Involved in the disassembly and reassembly of the Golgi before and after mitosis. | B1 |
